# GRAF1-dependent endocytotic processes and the Golgi apparatus contribute to novel intermediate stages of early ciliogenesis

**DOI:** 10.64898/2026.04.02.716088

**Authors:** Kerstin N. Schmidt, Korbinian Buerger, Olga Maier, Anita Zügner, Larissa Osten, Helga Othmen, Yulia Zaytseva, Anita Hecht, Reinhard Rachel, Ralph Witzgall

## Abstract

The intracellular cilia assembly pathway is a complex, multistep process that requires the continuous and coordinated incorporation of membrane material. However, how membrane remodeling occurs during early ciliogenesis is not yet understood. Moreover, the identity of the organelle(s) that supply membrane material for the nascent cilium has yet to be determined. Here, we extend the current model of primary cilia formation by showing that randomly attached distal appendage vesicles and tubules fuse laterally to generate a doughnut-shaped membrane structure. Centripetal fusion events follow to close the central hole. Our data demonstrate that both the Golgi apparatus and endocytotic pathways independently contribute to ciliogenesis. We identify the endocytotic protein GRAF1 as being essential during the early stages of ciliogenesis and for the delivery of plasma membrane-derived material to the developing ciliary membrane. Our three-dimensional ultrastructural analysis uncovers previously unrecognized intermediate stages in the intracellular cilia assembly pathway with GRAF1 as a novel regulator of ciliogenesis.

## Introduction

Over the recent years primary cilia have taken center stage when it comes to explaining the pathogenetic events underlying a number of hereditary diseases such as polycystic kidney diseases, Bardet-Biedl syndrome, retinal degeneration and skeletal dysplasias. Primary cilia are singular antenna-like extensions of most cell types in mammalian organisms that serve chemo-and mechanosensory functions and act as hubs for several signal transduction cascades. Two different pathways how primary cilia form have been described, the intracellular ^1,2^ and extracellular ^1,3^ pathways. This nomenclature, however, is somewhat misleading because in both pathways the primary cilium develops from the mother centriole in the cytoplasm, and even in the extracellular pathway no steps are known to take place on the extracellular side of a cell. Little is known about the extracellular pathway in which the mother centriole migrates up from its perinuclear position towards the plasma membrane where the axoneme grows out and pushes the plasma membrane outwards into the extracellular space. Most attention has been paid to the intracellular pathway where a primary cilium is pre-formed in the cytoplasm at the distal end of the mother centriole and subsequently fuses with the overlying plasma membrane to gain access to the extracellular space.

Transmission electron microscopic investigations published in 1962 still serve as an argument that the membrane material of the primary cilium derives from the Golgi apparatus ^2^, another publication almost ten years later pointed into the same direction ^4^. Strong data for the involvement of the Golgi apparatus in ciliogenesis are lacking, the underlying argument essentially refers to the close spatial connection between the centrosome and the Golgi apparatus. Since then, not much has changed of our view except that an earlier stage has been described which is characterized by the presence of individual vesicles at the distal appendages^5^. Accordingly, current thinking suggests that vesicles dock at the distal appendages of the mother centriole, then a primary ciliary vesicle forms and an axoneme starts to grow out, thereby pushing the center portion of the ciliary vesicle forward so that it assumes a cap- and bell-like shape (Fig. 1a).

**Figure 1.**
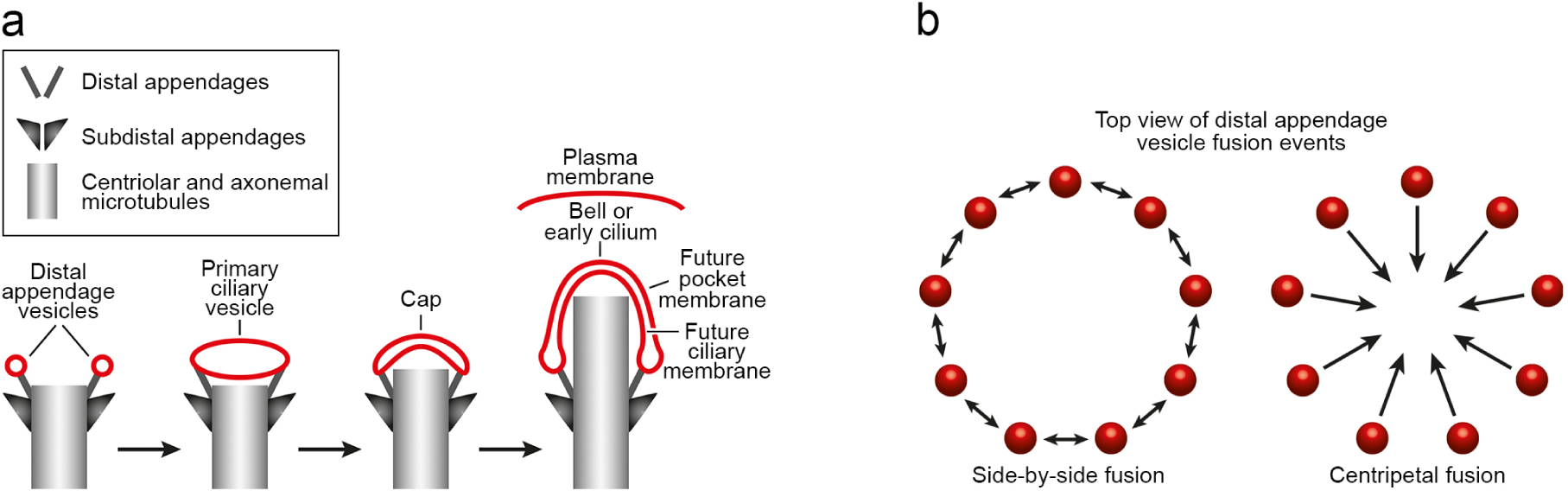
Cartoon highlighting distinct ultrastructural stages of primary ciliogenesis. (a) Conventional model for intracellular ciliogenesis. Vesicles dock at the distal appendages of the mother centriole and then fuse to form a primary ciliary vesicle which finally develops into the ciliary membrane. (b) Possible fusion routes of distal appendage vesicles to form a continuous membranous structure covering the distal end of the mother centriole. Membranes are depicted in red.

The picture, however, has become more complex because some publications implicate endocytotic processes in the formation of primary cilia. The simultaneous knock-down of Rab11a and Rab11b ^6^ and the knock-down of Rab11b ^7^ (the small monomerice GTPase Rab11 is associated with the recycling endosome ^8–10^) results in a lower number of ciliated cells. A negative effect on ciliogenesis is also observed upon the knock-down of the Arf GTPase ASAP1 ^11^ which interacts with Rab11 and regulates the trafficking of transferrin and its receptor^12^. Again by a knock-down approach it was demonstrated that the phosphotyrosine phosphatase PTPN23 ^11^ and EHD1 ^5^ participate in the development of primary cilia. Both PTPN23 ^13^ and EHD1 ^14^ are involved in the recycling of endocytosed proteins such as the transferrin receptor. A role of the recycling endosome in ciliogenesis is also supported by the knock-down of Ahi1, a protein mutated in patients suffering from Joubert syndrome. Ahi is important for the recycling of transferrin to the plasma membrane, its knock-down leads to an inhibition of ciliogenesis ^15^. Whereas ciliogenesis is reduced upon the knock-down of Rab11, ASAP1, PTPN23, EHD1 and Ahi1, the expression of a constitutively active Rab5 mutant blocks ciliogenesis ^16^. Rab5 is associated with the early endosome ^8–10^, it is noteworthy in this context that constitutively active Rab5 causes enlarged early endosomes and slows down the recycling of transferrin to the plasma membrane ^16,17^.

## Results

### STEM tomography reveals distal appendage tubules, the ‘doughnut’, bi-concave and hood stages of early ciliogenesis, and membrane fusion events

The consecutive stages of the intracellular cilia assembly pathway (Fig. 1a) have mainly been described based on the electron microscopical analysis of two-dimensional (serial) ultrathin sections (e.g. ^2,5,18–20^). However, a comprehensive three-dimensional characterization of ciliogenesis has not been carried out yet, in particular it is not known in what order distal appendage vesicles fuse with each other (Fig.1b) and where these vesicles originate from. We established a protocol for correlative light and electron microscopy (CLEM) to characterize membrane remodeling processes during early ciliogenesis. To this end, we generated a stably transfected human retinal pigment epithelial 1 (RPE1) cell line constitutively synthesizing a centrin1-EGFP fusion protein as a centrosomal marker ^21^ and an mCherry-Rab8a fusion protein to identify nascent primary cilia ^6^. RPE1 cells were serum-starved for 3 to 10 hours to induce ciliogenesis, and individual cells in which mCherry-Rab8a localized in a punctate pattern to one of the two centrin1-EGFP-positive centrioles were identified by fluorescence microscopy (Fig. 2a). Subsequently, these specific centrosomal regions were subjected to scanning transmission electron microscopy (STEM) tomography to obtain three-dimensional ultrastructural data with a high resolution ^22,23^. In a final step, the tilt series were three-dimensionally reconstructed using the IMOD software package ^24^.

**Figure 2.**
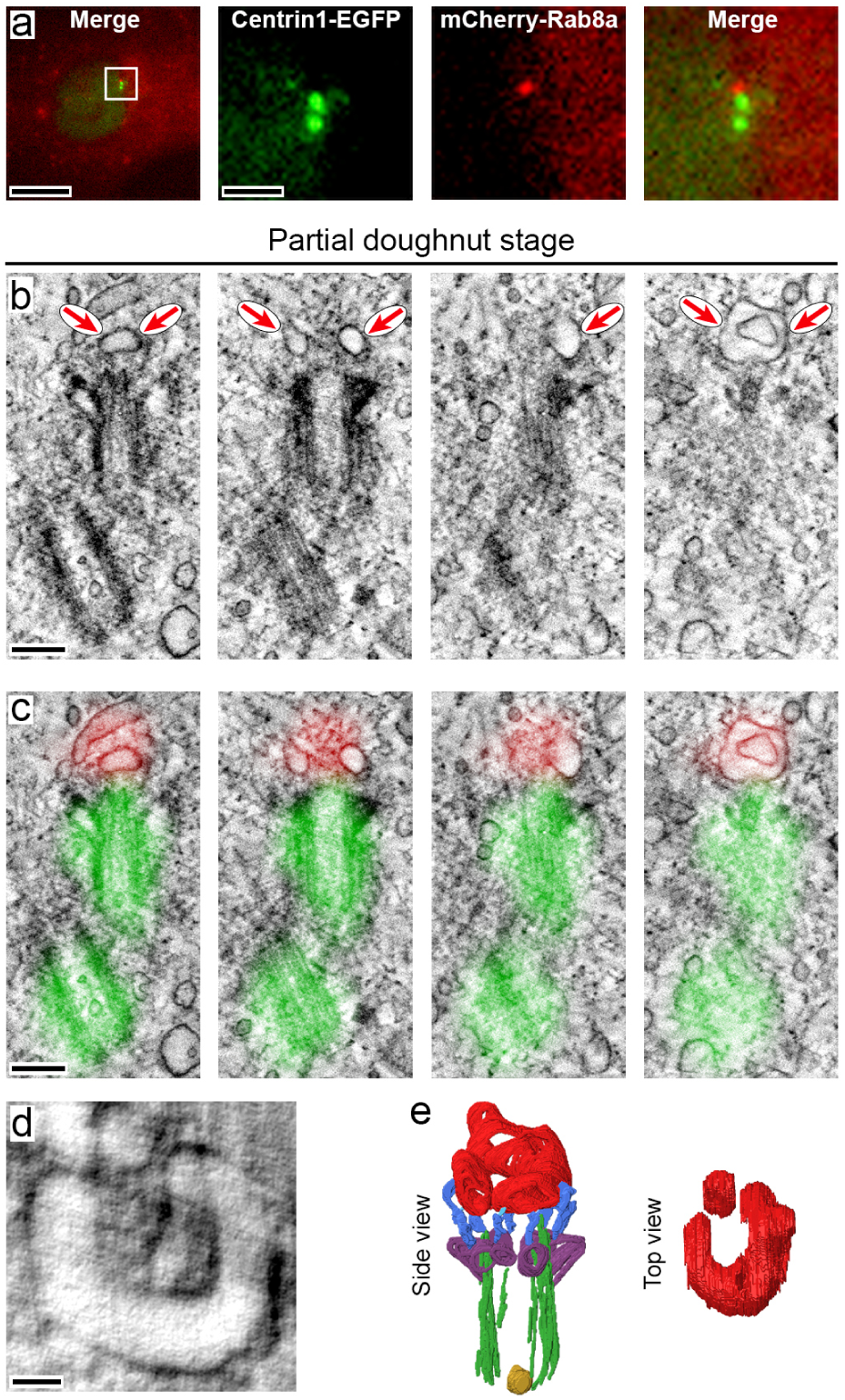
Identification of the partial doughnut stage of ciliogenesis by correlative light and electron microscopy. (a) Representative fluorescence image of a stably transfected RPE1 cell constitutively producing centrin1-EGFP and mCherry-Rab8a fusion proteins after 10 hours of serum starvation. It can be seen that Rab8a (red) accumulates at one of the centrioles (green) in a punctate fashion. The centrosomal region in the white box is shown at a higher magnification in the three panels on the right. (b, c) STEM (scanning transmission electron microscopy) tomography of the same position without (b) and with (c) an overlay of the fluorescence micrograph demonstrates the presence of the mCherry-Rab8a fusion protein at the distal end of the mother centriole (labeled by centrin1-EGFP). Red arrows point to the membrane of the nascent cilium. The slices lie ∼55 nm apart. (d, e) Three-dimensional reconstruction of the tomogram reveals the partial doughnut stage of cilium formation. A cross section of the nascent ciliary membrane structure is shown in panel d, the segmented nascent cilium in e. Microtubules, green; subdistal appendages, purple; distal appendages, dark blue; tether between distal appendages and nascent ciliary membrane, light blue; nascent ciliary membrane, red; intracentriolar vesicle, yellow. Bars: 10 µm (overview in a), 2 µm (higher magnification in a), 200 nm (b, c), 50 nm (d).

Centrosomal regions which had been documented before by fluorescence microscopic imaging were re-identified in STEM tomograms using the open source software package eC-CLEM ^25^. For each tomogram of a centrosomal region we confirmed the correlation of the centrin1-EGFP signals with the two centrioles as well as the overlap of the mCherry-Rab8a signal with the developing ciliary membrane structure at the distal end of the basal body (Fig. 2a-c). To facilitate the analysis of our three-dimensional data, centriolar and axonemal microtubules of the basal body, the subdistal and distal appendages with the associated membranous structures, and vesicles in the lumen of the basal body were segmented in selected tomograms. We routinely identified most, although sometimes not all, of the nine subdistal and distal appendages. Occasionally it was impossible to decide whether a given structure lay close enough to a distal appendage to be counted as attached. Views from different angles and contextual evidence such as a circular arrangement of vesicles proved helpful, but whenever no conclusive decision could be arrived at we opted for a more conservative approach and did not consider a membrane-lined structure as being associated with the mother centriole.

Our three-dimensional ultrastructural analysis revealed novel intermediate stages of cilia formation. The earliest and most frequent membrane structure attached to the distal appendages resembled an incomplete ring that we named the partial doughnut stage (13 out of 35 tomograms, i.e. 37%) (Fig. 2b-e). To be considered as a partial doughnut we required the incomplete ring to be attached to at least 5 of the 9 distal appendages, thereby covering more than half of the circumference. Thus, the earliest stage at which mCherry-Rab8a localizes to the nascent ciliary membrane compartment is the partial doughnut stage (see below, Fig. 4k). Another membrane structure that we detected less frequently resembled the shape of a disc with two opposite concavities, hence we named it the bi-concave stage of cilia formation (1 out of 35 tomograms, i.e. 3%) (Fig. 3a-c). Furthermore, we identified structures in which the future pocket membrane had extended already and the future ciliary membrane either ran perpendicular to the centriolar axis (Fig. 3d-f) or parallel to the future pocket membrane (suppl. Fig. 1). We named the first structure the hood stage (2 out of 35 tomograms, i.e. 6%) and the second structure the cap stage (1 out of 35 tomograms, i.e. 3%) of cilia formation. The latter is followed by the bell stage (5 out of 35 tomograms, i.e. 14%), at which time the axoneme had begun to grow out (Fig. 3g-j) ^26^. Finally, we also saw rather short primary cilia (13 out of 35 tomograms, i.e. 37%). Starting at the partial doughnut stage we detected tubular structures connected to the nascent and already established primary cilium (15 out of 35 tomograms, i.e. 43%) (e. g. suppl. Fig. 2) similar to what has been described before ^27^. Some of these tubules ended in a blind fashion in the cytoplasm but in most cases it was not possible to follow them to their very end to decide whether they were connected to the plasma membrane or not.

**Figure 3.**
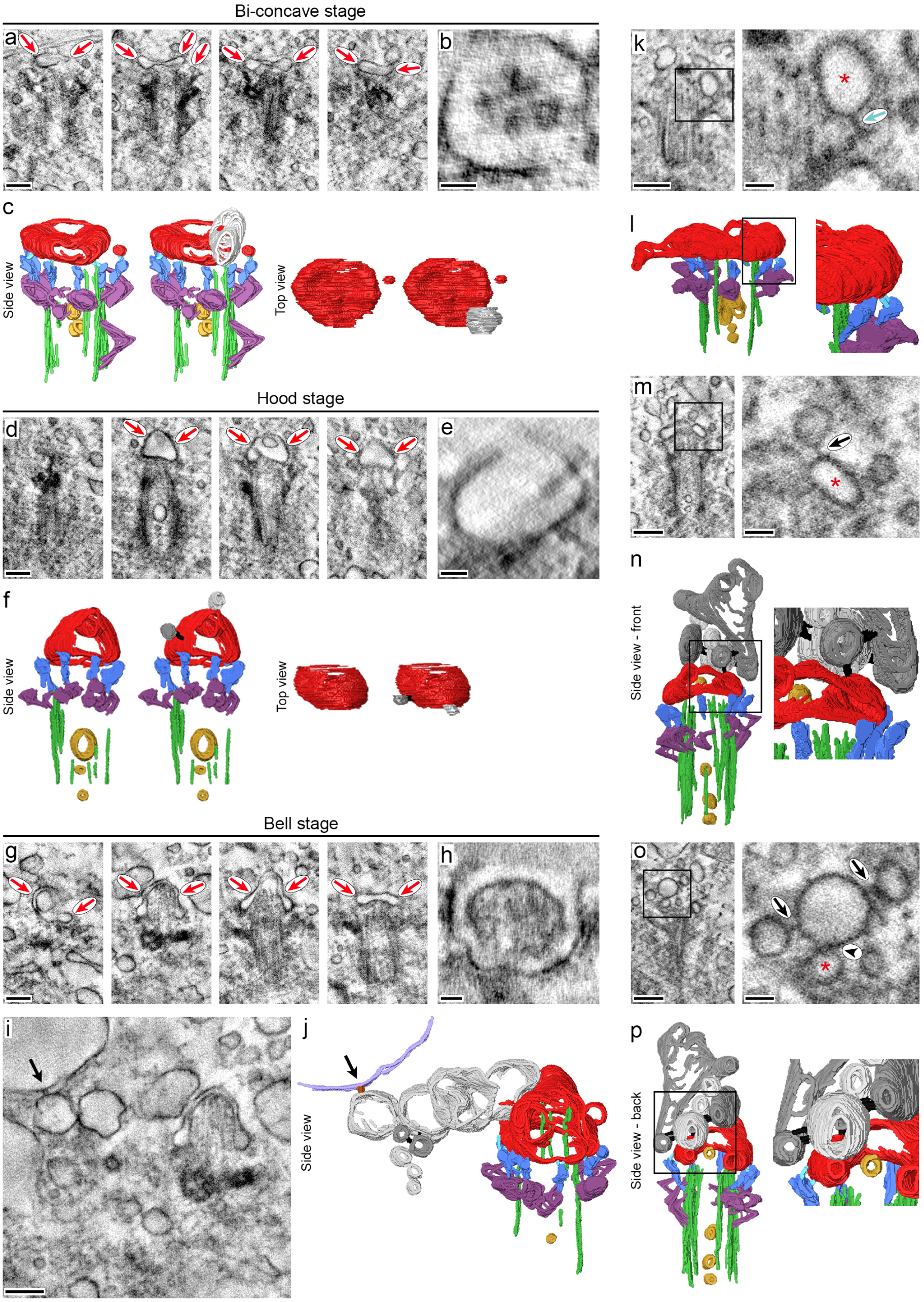
Discovery of the bi-concave, hood and bell stages of cilium formation by correlative light and electron microscopy. Tomograms of RPE1 cells in which mCherry-Rab8a accumulated at the centrin1-EGFP-postive centrosome (a-j, m-p) and of a RPE1 cell expressing centrin1-EGFP and mCherry (k, l). All cells were serum starved for 10 hours. (a-j) Three tomograms demonstrating distinct stages of ciliogenesis. Shown in each case are four slices lying ∼55 nm apart (a, d, g), a cross section of the nascent ciliary membrane structure (b, e, h) and the segmented nascent cilium without (segmentations on the left in c and f) and with (segmentations on the right in c and f, as well as j) incoming vesicles. In panel (i) a larger region between the mother centriole and the plasma membrane is shown to demonstrate the connection of the leftmost vesicle to the plasma membrane (lilac) via a tether (brown structure highlighted by an arrow). Red arrows in the tomograms point to the membrane of the nascent cilium. Vesicles in dark grey are attached to other membranes by tethers (shown in black), vesicles in light grey directly touch other membranes. (k, l) Selected plane of a tomogram and corresponding segmentation of the nascent cilium demonstrating a tether (light blue arrow) between a distal appendage and the nascent cilium (red asterisk). (m-p) Selected planes of a tomogram and corresponding segmentation of the nascent cilium demonstrating membranes connected by tethers, and membranes directly touching each other (front view in m and n, back view in o and p). Arrows point to tethers, the arrow head points to membranes directly touching each other, red asterisks mark the nascent cilium. Overviews are shown on the left, higher magnifications of the boxed areas on the right (k-p). Microtubules, green; subdistal appendages, purple; distal appendages, dark blue; tethers between distal appendages and nascent ciliary membrane, light blue; nascent ciliary membrane, red; intracentriolar vesicle, yellow. Bars: 200 nm (a, d, g, i, overviews in k, m, o), 50 nm (b, e, h, higher magnifications in k, m, o).

**Figure 4.**
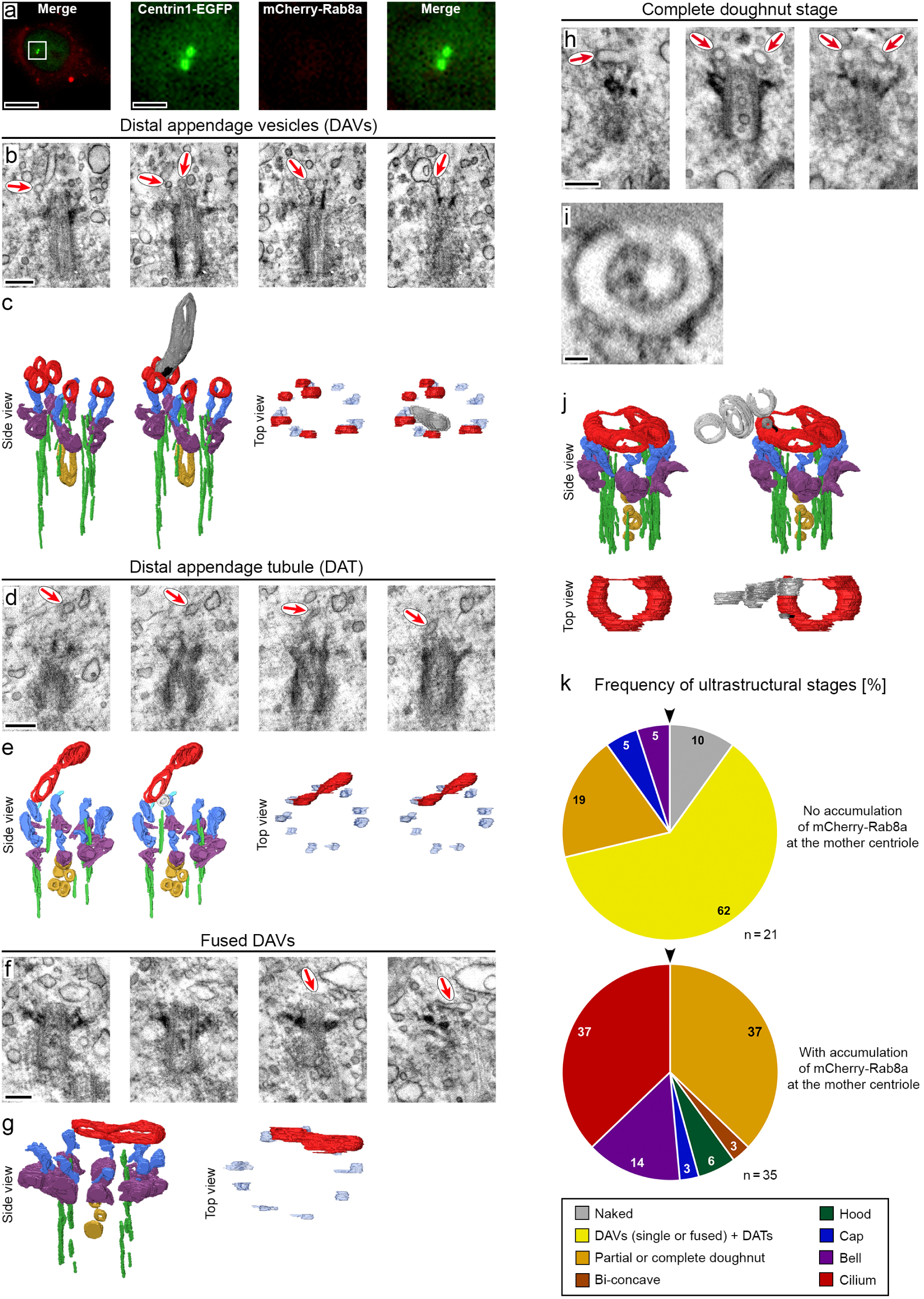
Rab8a acts preferentially after the distal appendage vesicle/tubule stage. Correlative light and electron microscopy of stably transfected RPE1 cells constitutively producing centrin1-EGFP and mCherry-Rab8a fusion proteins in which Rab8a has not accumulated at the centrosome. All cells were serum starved for 10 hours. (a) Representative fluorescence image of a cell. The centrosomal region in the white box is shown at a higher magnification in the three panels on the right. (b-g) The tomograms demonstrate the association of vesicles (b, c), tubules (d, e) and fused vesicles (f, g) with distal appendages. Shown in each case are four slices lying ∼80 nm (two leftmost slices) and ∼27 nm (slices two to four) (b), ∼27 nm (d) and ∼58 nm (f) apart, and the segmented nascent cilia without (left) and with (right) incoming vesicles (c, e). The dark grey tubular structure is tethered to one of the distal appendage vesicles (c), the light grey vesicular structure directly touches a distal appendage tubule (e). Red arrows in the tomograms point to the membrane of the nascent cilium. (h-j) Tomogram illustrating a complete doughnut-shaped membrane structure. Shown are three slices lying ∼80 nm apart (h), a cross section of the nascent ciliary membrane structure (i) and the segmented nascent cilium without (left) and with (right) incoming vesicles (j). Red arrows in the tomograms point to the doughnut-shaped membrane structure. The vesicle in dark grey is attached to the doughnut-shaped membrane structure by a tether (shown in black), vesicles in light grey directly touch the doughnut-shaped membrane structure and each other. (k) Pie charts depicting the frequency (in %) of ciliogenesis stages in cells where the mCherry-Rab8a fusion protein did not (top) and did (bottom) accumulate at the centrosome. Data include experiments with cells which were serum-starved for 3, 3.5 and 10 hours. Early to late stages are portrayed clockwise starting at the arrow head. *n* indicates the number of tomograms out of 6 (upper pie chart) and 14 (lower pie chart) independent experiments. Microtubules, green; subdistal appendages, purple; distal appendages, dark blue; tether between distal appendage and nascent ciliary membrane, light blue; nascent ciliary membrane, red; intracentriolar vesicles, yellow. Bars: 10 µm (overview in a), 2 µm (higher magnification in a), 200 nm (b, d, f, h), 50 nm (i).

We routinely found many vesicles in the vicinity of the mother centriole (Fig. 3, 4). Upon close examination filamentous structures, which had been missed in the past, apparently tethered some of those vesicles and also the membrane structure of the nascent primary cilium to the distal appendages (arrow in Fig. 3k, l). Moreover, tethering filaments were also observed between membranes which had already attached at the distal appendages and newly arriving vesicles (arrows in Fig. 3m-p). In addition to tethered vesicles we identified vesicles that lay immediately adjacent to the nascent primary ciliary membrane with both membranes touching each other (arrow head in Fig. 3o). Such arrangements were found in 27 out of 35 (77%) tomograms. Sometimes we noted additional vesicles tandemly connected to the first one like a string of pearls (Fig. 3g-j, m-p). In one instance this chain of vesicles was also tethered to the plasma membrane (arrows in Fig. 3i, j). On the other hand, we also observed structures obviously budding out from the future ciliary pocket membrane into the cytoplasm at various stages of ciliogenesis. Some of them were covered by electron-dense material which formed spikes very much resembling clathrin coats (3 out of 35 tomograms, i.e. 9%) (green arrow heads in suppl. Fig. 1). While clathrin-coated pits have been repeatedly observed to bud from the mature ciliary pocket membrane ^1,28^, our observation for the developing ciliary membrane compartment is novel.

To analyze events earlier than the partial doughnut stage and which do not depend on Rab8a we utilized the same cell line and turned to cells where Rab8a was present in the cytoplasm but had not been recruited to the centrosome yet (Fig. 4a). To exclude primary cilia undergoing disassembly we made sure by fluorescence microscopy and electron tomography that the centrosomes under investigation did not contain more than 2 centrioles as would be expected after cells had entered the S-phase of the cell cycle. In contrast to cells in which Rab8a had accumulated at the centrosome, we mainly identified stages earlier than the partial doughnut stage such as naked mother centrioles (2 out of 21 tomograms, i.e. 10%), and basal bodies with one or several vesicles attached to distal appendages (Fig. 4b, c) (13 out of 21 tomograms, i.e. 62%). Surprisingly, in addition to distal appendage vesicles we identified tubular membrane structures associated with the distal appendages that we named distal appendage tubules (Fig. 4d, e). Distal appendage vesicles and tubules were never observed in cells where mCherry-Rab8a had accumulated at the mother centriole (Fig. 4k) confirming earlier observations that Rab8a regulates ciliogenesis at more advanced stages ^5^. In none of the analyzed centrosomal regions were all distal appendages occupied by vesicles or tubules at the same time. Therefore fusion events between adjacent distal appendage vesicles and tubules occurred already before all distal appendages were occupied (Fig. 4f, g). These fused distal appendage vesicles were categorized as partial doughnut-shaped membrane structures only when they were attached to more than 4 distal appendages (4 out of 21 tomograms, i.e. 19%). Furthermore we identified one complete ring-shaped membrane structure in this setup and thus a closed doughnut stage (Fig. 4h-j). Although we detected the cap and bell stages of cilia formation (1 each out of 21 tomograms, i.e. 5%), no outgrowing cilia were found in cells in which Rab8a had not yet been recruited to the mother centriole (Fig. 4k).

To exclude the possibility that the presence of (partial and complete) doughnut-shaped membrane structures and of distal appendage tubules resulted from the overexpression of mCherry-Rab8a we also investigated centrosomes of RPE1 cells that stably synthesized centrin1-EGFP and transiently expressed mCherry (suppl. Fig. 3). Again distal appendage tubules were identified, and once more the most frequent stage was the partial doughnut stage (9 out of 25 tomograms, i.e. 36%) (suppl. Fig. 3). Newly arriving membrane material connected to the membrane of the nascent cilium by tethers or directly touching it was present in 63% (15 out of 24 tomograms) of cells producing mCherry and in 77% (27 out of 35 tomograms) of cells in which mCherry-Rab8a had accumulated at the mother centriole. We observed tubular structures connected to the nascent cilium in 47% (8 out of 17 tomograms) of cells producing mCherry and in 43% (15 out of 35 tomograms) of cells in which mCherry-Rab8a had accumulated at the mother centriole. Furthermore, clathrin-coated pits were detected in 8% (2 out of 24 tomograms) of cells producing mCherry and in 9% (3 out of 35 tomograms) of cells where mCherry-Rab8a had accumulated at the mother centriole. Hence, the novel intermediate stages of ciliogenesis, the arrival of additional membrane material at the nascent cilium, the formation of tubules and the budding of clathrin-coated pits at the nascent cilium are general features of the intracellular cilia assembly pathway and form independently of the fact whether mCherry-Rab8a is overexpressed or not.

### Characterization of intracentriolar vesicles

A regular finding in our tomograms was the presence of intracentriolar vesicles both in the mother and the daughter centriole (e.g. Fig. 2b, Fig. 3a, d, m, suppl. Fig. 4a-d) similar to what has been described before (e.g. in ^2,29^). Their function is not known, they may constitute a population of incoming or outgoing vesicles or both. If these vesicles represented mostly incoming vesicles, i.e. vesicles contributing to ciliogenesis, one would expect a larger number in the mother centriole than in the daughter centriole because only the mother centriole will give rise to a primary cilium. Although on average more vesicles were observed inside the mother centriole, this difference did not reach statistical significance (suppl. Fig. 4e). As Rab8a promotes ciliogenesis ^30^ more or larger intracentriolar vesicles might be expected in the presence of the mCherry-Rab8a fusion protein. This turned out not to be the case (suppl. Fig. 4e, f). Finally, we explored the relationship between the number of intracentriolar vesicles in the mother and the daughter centriole, wondering whether a larger number of vesicles in the mother centriole correlated with a lower number in the daughter centriole and vice versa. No obvious pattern emerged (suppl. Fig. 4g, h) and it therefore rather seems that the number of vesicles in the mother and the daughter centrioles is independently regulated. Moreover, we never observed fusion events between intracentriolar vesicles and the developing ciliary membrane compartment attached to the distal appendages of the basal body. In summary, the function of the intracentriolar vesicles during ciliogenesis remains obscure.

### Endocytosed membrane material is delivered to the nascent ciliary membrane compartment

For the primary cilium to form, a steady delivery of vesicles has to be guaranteed. Over the years scarce evidence has been published that the endosomal compartment, in particular the recycling endosome, contributes to the formation of primary cilia ^6,11,31–33^. In *Chlamydomonas,* Arp2/3 complex-dependent endocytosis delivers ciliary membrane proteins to the growing flagellum ^34^. To investigate whether endocytosed membrane material is directed to the developing ciliary membrane compartment during the intracellular cilia assembly pathway we incubated RPE1 cells with horseradish peroxidase (HRP)-conjugated wheat germ agglutinin (WGA). The lectin WGA binds to widespread sugar residues of membrane proteins such as *N*-acetyl-*D*-glucosamine ^35,36^ and sialic acid residues ^37,38^. Since as a protein WGA cannot permeate membranes, it will only recognize the extracellular portion of cell surface glycoproteins. A subsequent histochemical reaction for HRP and incubation with OsO_4_ leads to the formation of an osmiophilic and therefore electron-dense diaminobenzidine precipitate at the sites where WGA has bound, which allows the visualization of endocytosed plasma membrane material. RPE1 cells stably expressing centrin1-EGFP were serum-starved for 4 hours to initiate cilium formation and cultured for the last 15 minutes at 37°C in the presence of the WGA-HRP conjugate. A short incubation time was chosen to avoid retrograde trafficking of WGA-HRP to the Golgi apparatus ^39–41^ which would make it impossible to discriminate between endocytotic routes and the secretory pathway. Electron-dense diaminobenzidine precipitates were detected by transmission electron microscopy of 80 nm thick sections at the plasma membrane and at cytoplasmic vesicular structures (suppl. Fig. 5). The specificity of our approach was corroborated by the finding that many membrane-lined structures in the cytoplasm were not labeled (suppl. Fig. 5). As an additional control, we could not detect any electron-dense material at the plasma membrane and in intracellular vesicles when the histochemical reaction was carried out with cells which were incubated in the absence of WGA-HRP (suppl. Fig. 5). To examine whether endocytosed membrane material is incorporated into the nascent ciliary membrane, centrosomal regions of RPE1 (centrin1-EGFP) cells incubated in the presence of WGA-HRP were identified in the fluorescence microscope and then subjected to electron tomography. In 25 out of 42 cells (i.e. 60%) nascent primary cilia, from the distal appendage vesicle/tubule stage to the bell stage, were labeled with electron-dense material (Fig. 5a-f). No labeling of the nascent ciliary membrane was detected when the cells were incubated in the absence of WGA-HRP (Fig. 5g, h). In principle, WGA-HRP might reach the nascent primary cilium not only by endocytosis but also via tubular extensions that connect the developing ciliary membrane compartment with the plasma membrane and thus with the extracellular space (suppl. Fig. 2 and ^27^). Careful three-dimensional analysis yielded 12 labeled nascent primary cilia that were completely obtained in the tomogram and either had not elaborated tubular extensions or the tubular extensions ended blindly in the cytoplasm without any connection to the plasma membrane. Therefore we conclude that endocytosed membrane material contributes to the growing ciliary membrane compartment. As expected, most of the fully formed primary cilia (24 out of 27, i.e. 89%) were routinely labeled with WGA-HRP (Fig. 5i) because they either had direct contact to the extracellular space or were connected to the plasma membrane by tubules.

**Figure 5.**
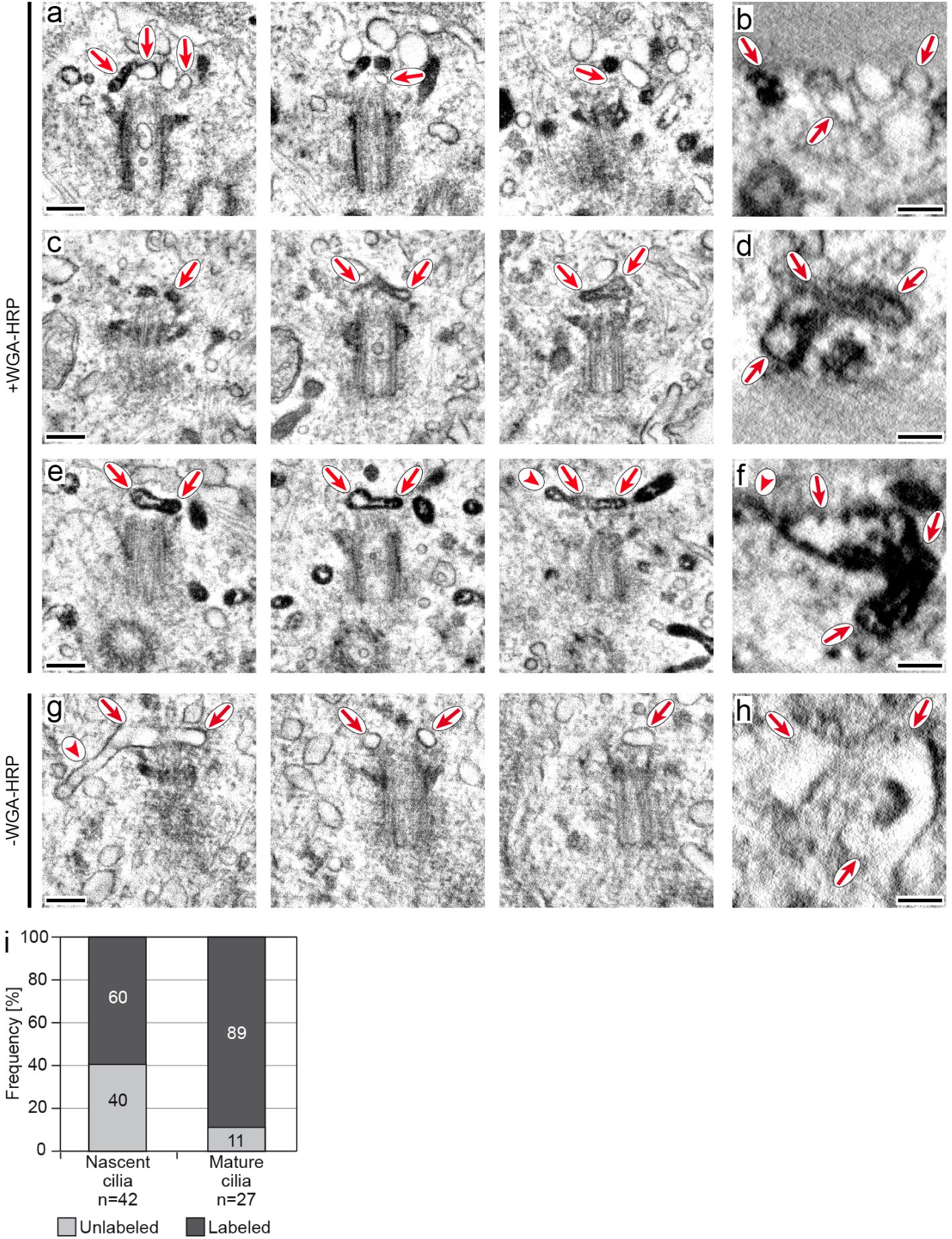
Endocytosed membranes associate with the mother centriole. (a-f) Tomograms of RPE1 cells producing a centrin1-EGFP fusion protein. Cells were serum-starved for 4 hours and exposed to a fusion protein between wheat germ agglutinin (WGA) and horseradish peroxidase (HRP) for the last 15 minutes of serum starvation. (g, h) RPE1 cells which were not exposed to WGA-HRP served as a negative control. Red arrows point to the membrane of the nascent cilium, red arrow heads to tubular extensions. Shown in each case are three slices lying ∼46 nm apart (a), ∼55 and ∼80 nm nm apart (c), ∼55 nm apart (e), ∼55 and ∼110 nm apart (g). Respective cross sections of the nascent ciliary membrane structure are seen in b, d, f, h. (i) Quantification of nascent and mature cilia labeled with WGA-HRP in cells treated as described above. Over half of the nascent cilia and almost all the mature cilia were labeled by WGA-HRP. *n*, number of tomograms out of 2 independent experiments. Bars: 200 nm (a, c, e, g), 100 nm (b, d, f, h).

### Clathrin- and caveolin-dependent endocytosis are dispensable for cilia formation

In order to identify the endocytosis routes that contribute membrane material to ciliogenesis, we first examined clathrin- and caveolae-dependent endocytosis as the most common endocytosis pathways ^42,43^. Here, the scission of invaginating vesicles is mediated in large part by dynamins, a family of monomeric high-molecular weight GTPases ^44^. The incubation with dynasore, a chemical inhibitor of dynamins ^45^, resulted in a statistically significant decrease in the number of ciliated cells after 24 hours of serum deprivation, which may be due to a slowly developing toxic effect of dynasore, because the effect was rather the opposite (without reaching statistical significance) at earlier time points (Fig. 6a). To investigate the effect of dynamin more specifically, RPE1 cells were transiently transfected with an expression plasmid encoding the dominant-negative K44A mutant of dynamin-2 ^46^. Compared to EGFP-expressing cells, a statistically significant increase in the number of ciliated cells was observed at the beginning and at 24 hours of serum withdrawal but not between those time points (Fig. 6b). To address clathrin-mediated endocytosis directly we transiently expressed dominant-negative forms of EPS15 [EPS15 (DIII)] ^47^ and AP180 (AP180-C) ^48^, two proteins involved in the formation of the clathrin cage ^42^. A statistically significant increase in the number of ciliated cells was detected for the dominant-negative mutant of EPS15 but not for AP180 (Fig. 6b). The increase in ciliated cells upon dynamin (K44A)-EGFP and EGFP-EPS15 (DIII) expression was confirmed in a time-course with asynchronously growing RPE1 cells (i.e. cells cultured in the presence of serum) at 48 and 24 hours after transfection, respectively (Fig. 6c). These data argue against an essential function of clathrin-mediated endocytosis in the formation of primary cilia, which is in line with a previous report ^1^. If anything, we noticed more cilia when clathrin-mediated endocytosis was inhibited. The treatment of cells with methyl-β-cyclodextrin depletes the plasma membrane of cholesterol which is necessary for caveolae-mediated endocytosis ^49,50^. Neither incubation with methyl-β-cyclodextrin (Fig. 6d) nor the transient transfection of expression plasmids encoding dominant-negative mutants of caveolin-1 (ΔN1-81 ^51^ and P132L ^52^) (Fig. 6e) inhibited ciliogenesis compared to control-treated cells. Accordingly our results argue that neither clathrin- nor caveolin-mediated endocytosis contribute to cilia formation.

**Figure 6.**
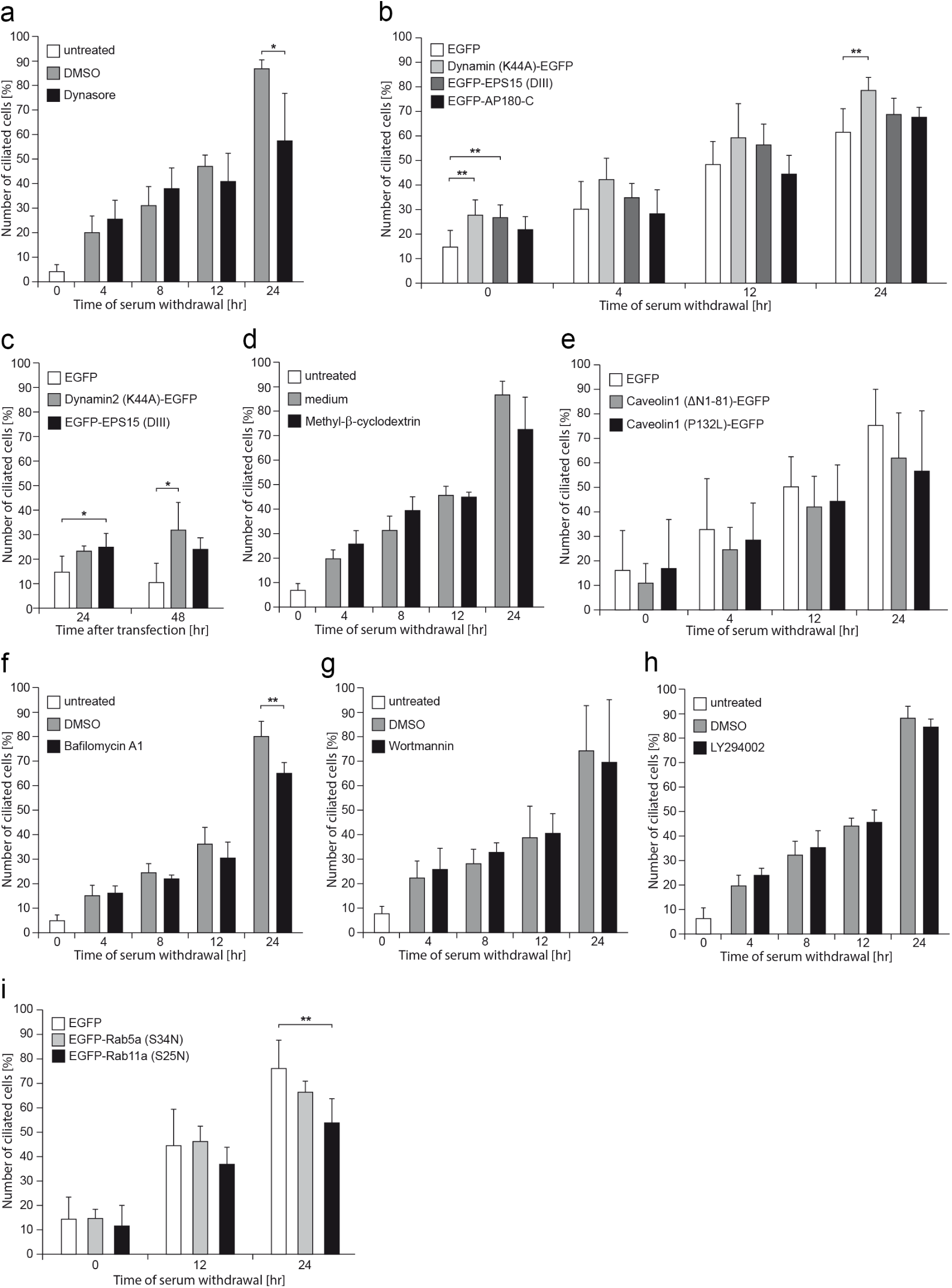
Conventional forms of endocytosis do not contribute to the formation of primary cilia. RPE1 cells were serum-starved and treated with various small molecules (dynasore, methyl-β-cyclodextrin, bafilomycin A1, wortmannin, LY294002) for the indicated periods of time (a, d, f, g, h). When RPE1 cells were transfected with expression plasmids encoding the respective dominant-negative mutant proteins (b, c, e, i), serum starvation was started 24 hours after transfection and carried out for the indicated periods of time. In the former case the solvent DMSO (except in d where medium was used as negative control), in the latter case the expression plasmid for EGFP served as the negative controls. Shown are the mean + standard deviation, *n* = 4-7 independent experiments. *, *p* < 0.05; **, *p* < 0.01.

### Inhibition of different endosomal compartments does not impair ciliogenesis

The dispensability of clathrin- and caveolae-mediated endocytosis for ciliogenesis led us to inhibit different endosomal compartments ^8^. Bafilomycin A1 inhibits a vacuolar H^+^-ATPase which is responsible for the increasing acidification along the endosomal pathway ^53^. Furthermore, the maturation from the early to the late endosome is believed to depend on the action of phosphatidylinositol (3) kinases ^54^ which can be blocked by wortmannin ^55,56^ and the non-selective inhibitor LY294002 ^57,58^. If the vesicles destined to contribute to ciliogenesis are derived from the late endosome or an even more downstream compartment, then the incubation with these substances should inhibit ciliogenesis. A statistically significant negative effect was only observed at 24 hours of treatment with bafilomycin A1 which, similar to what we observed for dynasore, may be due to a toxic effect of bafilomycin A1 (Fig. 6f). In the case of wortmannin and LY294002 no effect on ciliogenesis was noticed (Fig. 6g, h). Individual endosomal compartments were addressed more specifically by transfecting RPE1 cells with expression plasmids encoding dominant-negative mutants of Rab5a and Rab11a, monomeric small GTPases associated with early endosomes and recycling endosomes, respectively ^8,10^. Whereas expression of dominant-negative Rab5a had no effect on ciliogenesis, the expression of dominant-negative Rab11a reduced the number of primary cilia (Fig. 6i) which indicates that the recycling endosome is involved in ciliogenesis as has been suggested before ^6,16,32^.

### GRAF1 as a member of the CLIC/GEEC endocytotic pathway is involved in ciliogenesis

Since (apart from the recycling endosome) none of the conventional endocytosis pathways contributed to cilia formation, we turned our attention to alternative endocytotic routes. In addition to caveolae-mediated endocytosis, several other forms of endocytosis are subsumed into the category of clathrin-independent endocytosis ^59,60^. The scission of tubular- and ring-like structures formed at the plasma membrane within the CLIC (<u>cl</u>athrin-independent <u>c</u>arrier) endocytosis pathway proceeds without coats and dynamin ^61,62^, and the carriers fuse independently of Rab5 to build the GEEC [<u>g</u>lycosylphosphatidylinositol-anchored proteins (GPI-AP) <u>e</u>nriched early <u>e</u>ndosomal <u>c</u>ompartment] ^62,63^. Both features are consistent with the observations described above which led us to investigate whether the CLIC/GEEC pathway is involved in ciliogenesis. This endocytosis pathway depends on the multidomain, non-cargo protein GRAF1 (<u>G</u>TPase regulator <u>a</u>ssociated with focal adhesion kinase 1) ^63,64^. When GRAF1 was depleted in RPE1 cells by RNA interference a decrease in the number of ciliated cells was observed (Fig. 7a, b). The specificity of the siRNA was demonstrated by a rescue experiment in which ciliogenesis was partially restored upon transient expression of an RNAi-resistant EGFP-GRAF1 (rescue) expression construct (Fig. 7c, d).

**Figure 7.**
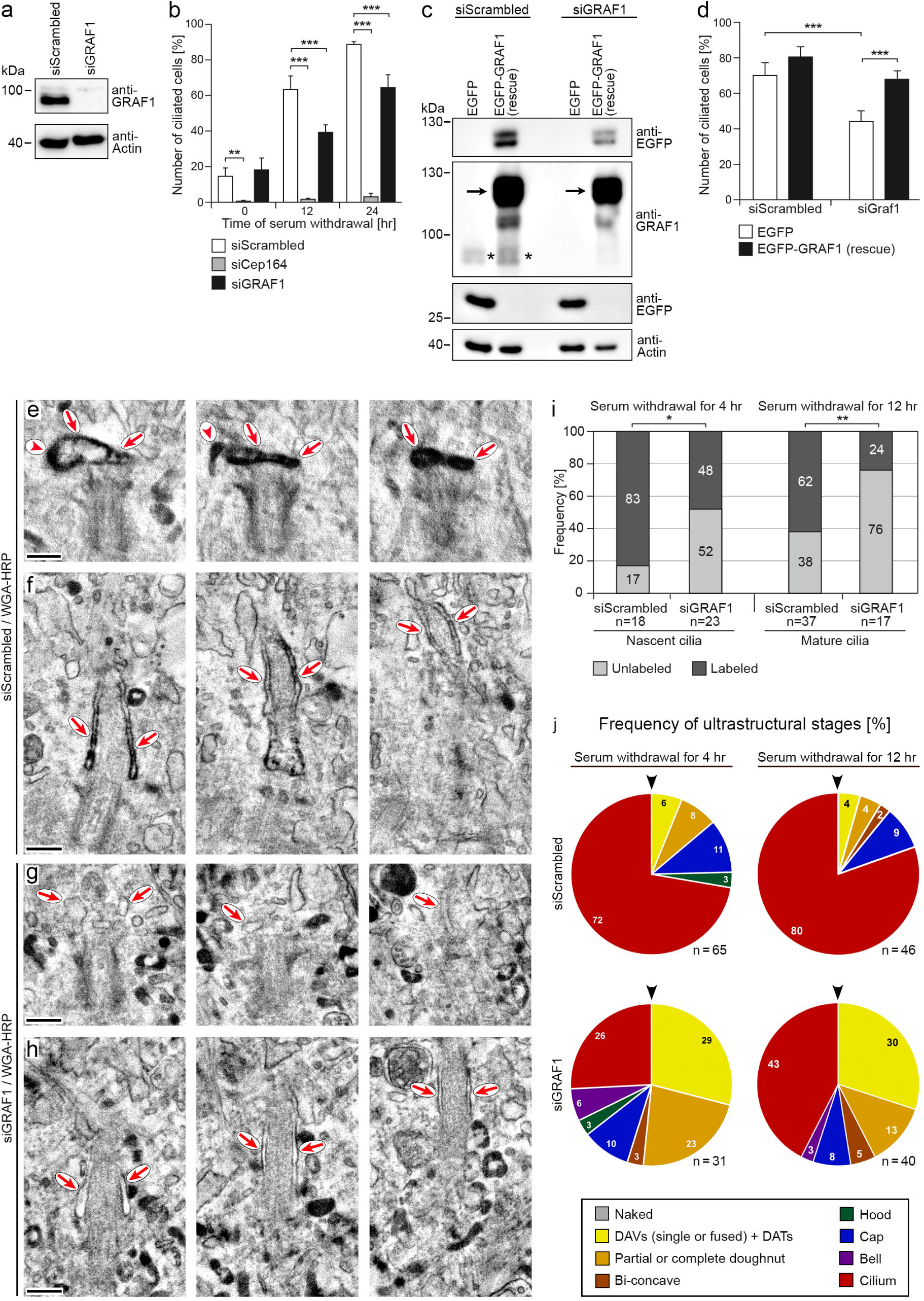
GRAF1 contributes to the formation of primary cilia. (a) A Western blot of RPE1 cells treated with the respective siRNAs for 48 hours demonstrates the efficient knock-down of GRAF1. Actin served as a loading control. (b) RPE1 cells were treated as described for panel a and serum-deprived for the indicated periods of time. The knock-down of GRAF1 leads to a reduced number of ciliated RPE1 cells after serum withdrawal. An siRNA against Cep164 served as a positive control. *n* = 5 independent experiments. (c, d) Rescue experiment with an expression plasmid for an siRNA-resistant EGFP-GRAF1 (rescue) mRNA. RPE1 cells were transiently transfected with the indicated expression plasmids. Twenty-four hours after the transfection they were subjected to treatment with the relevant siRNAs, another 24 hours later the cells were serum-starved for a further 24 hours. A Western blot demonstrates the successful depletion of the endogenous GRAF1 protein (asterisks) and expression of the exogenous fusion protein (arrows). The expression of EGFP served as a negative control (c). The EGFP-GRAF1 (rescue) fusion protein is able to restore ciliogenesis upon the knock-down with the siRNA against the GRAF1 mRNA. *n* = 6 independent experiments. (e-j) RPE1 cells producing centrin1-EGFP were treated with a scrambled siRNA or an siRNA against GRAF1. Forty-eight hours later the cells were serum-starved for 4 or 12 hours before they were exposed to a fusion protein between wheat germ agglutinin (WGA) and horseradish peroxidase (HRP) for 15 minutes. Three planes of tomograms of cells starved for 12 hours lying ∼80 nm (e), ∼72 nm (f), ∼70 nm (g) and ∼72 nm (h) apart are shown for each tomogram. Red arrows point to the membrane of the nascent cilium, arrow heads to tubular extensions (e-h). The bar graph (i) demonstrates that both nascent and mature cilia were labeled to a lower degree in GRAF1-depleted cells. (j) Pie charts depicting the frequency (in %) of ciliogenesis stages, early to late stages are portrayed clockwise starting at the arrow head. A knock-down of GRAF1 retards the development of primary cilia. *n* indicates the number of tomograms. *, *p* < 0.05; **, *p* < 0.01; ***, *p* < 0.001. Bars: 200 nm.

To identify the stage(s) of cilia formation that require GRAF1, RPE1 cells stably expressing centrin1-EGFP were depleted of GRAF1 and ultrastructurally analyzed by STEM tomography. At the same time, the cells were incubated with WGA-HRP to verify whether endocytosed membrane material is still delivered to the nascent primary cilium in the absence of CLICs. Cells were serum-deprived for 4 and 12 hours to assess nascent as well as mature cilia. Upon depletion of GRAF1, less RPE1 cells with labeled nascent and mature ciliary membranes were identified compared to control-depleted cells (Fig. 7e-i). Furthermore, while most of the tomograms from centrosomal regions of control-depleted cells showed an outgrown cilium after 4 hours (47 out of 65 tomograms, i.e. 72%) and 12 hours of serum starvation (37 out of 46 tomograms, i.e. 80%), a substantial number of GRAF1-depleted cells had only reached the stage of distal appendage vesicles and tubules at 4 hours (9 out of 31 tomograms, i.e. 29%) and 12 hours (12 out of 40 tomograms, i.e. 30%) of serum starvation (Fig. 7j).

### Involvement of the Golgi apparatus in the formation of primary cilia

Another organelle that is regularly brought up as a source for ciliary membrane material is the Golgi apparatus (for review see ^65–67^). Our regular finding of the Golgi apparatus in close vicinity to the centrosome is in line with long-standing observations published several decades ago. Ultrastructural investigations particularly from the 1960s are routinely quoted as evidence that the ciliary membrane material is transported from the Golgi complex to the centrosome, remarkably the only argument being their close spatial arrangement ^2,29,68^. However, direct experimental data for such an assumption have not been presented so far. We addressed this question by disrupting or inhibiting the Golgi apparatus by different means. First, we incubated RPE1 cells with brefeldin A (BFA), a fungal metabolite which inhibits guanine nucleotide exchange factors of the ARF family ^69,70^ thus resulting in the collapse of the Golgi apparatus ^71,72^. This led to a lower number of ciliated cells (Fig. 8a). The involvement of a functional Golgi apparatus was confirmed by treating RPE1 cells with two other chemical inhibitors of the Golgi apparatus, the ionophore monensin ^73^ and 30N12 ^74^. After 12 and 24 hours of serum starvation both compounds also produced a significant inhibitory effect on ciliogenesis (Fig. 8b, c). These experiments support the notion that the Golgi apparatus contributes to ciliogenesis but it is not known at what stage(s).

**Figure 8.**
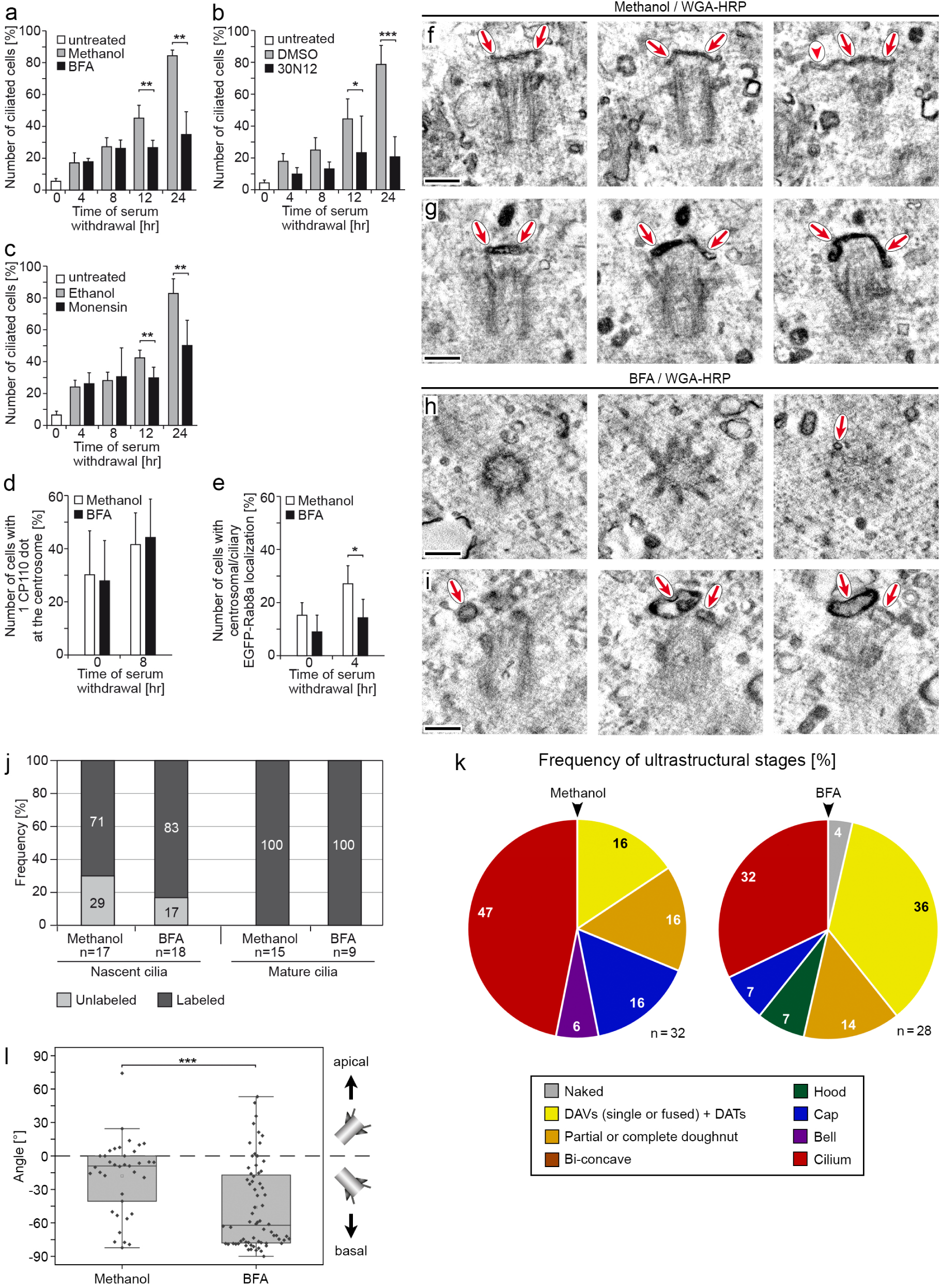
Involvement of the Golgi apparatus in the formation of primary cilia. (a-c) RPE1 cells were serum-starved and treated with the substances brefeldin A (BFA), 30N12 or monensin for the indicated periods of time. The number of ciliated cells was reduced significantly compared to the respective solvents methanol, DMSO and ethanol. *n* = 4-8 independent experiments. (d) RPE1 cells stably producing centrin1-EGFP were incubated for 2 hours with methanol and BFA in the presence of serum and an additional 8 hours in its absence. Treatment with brefeldin A had no effect on the abundance of cells with only a single centrosomal dot of CP110. (e) RPE1 cells stably producing EGFP-Rab8a were incubated for 2 hours with methanol or BFA in the presence of serum and an additional 4 hours in its absence. The number of cells in which Rab8a accumulated at the centrosome was reduced upon BFA treatment. (f-l) RPE1 cells stably producing centrin1-EGFP were incubated for 2 hours with methanol or BFA in the presence of serum and an additional 4 hours in its absence. For the last 15 minutes, the cells were exposed to a fusion protein between wheat germ agglutinin (WGA) and horseradish peroxidase (HRP). Three planes lying ∼80 nm (f), ∼60 nm (g), ∼55 nm (h) and ∼60 nm (i) apart are shown for each tomogram, red arrows point to the membrane of the nascent cilium, the arrow head to a tubular extension (f). The bar graph (j) demonstrates that nascent and mature cilia were labeled to the same degree both in methanol- and brefeldin A-treated cells. *n* indicates the number of tomograms. (k) Pie charts depicting the frequency (in %) of ciliogenesis stages, early to late stages are portrayed clockwise starting at the arrow head. Treatment of RPE1 cells with BFA retards the development of primary cilia. *n* indicates the number of tomograms. (l) Orientation of the mother centriole in RPE1 cells. Positive angles indicate an orientation of the distal end of the mother centriole towards the apical cell membrane, negative angles towards the basal cell membrane. The distal end of the mother centriole points towards the basal cell membrane after treatment with brefeldin A. *n* = 37 (methanol-treated cells) and 68 (BFA-treated cells). *, *p* < 0.05; **, *p* < 0.01; ***, *p* < 0.001. Bars: 200 nm.

Cep97 and CP110 are located at the distal end of both the mother and daughter centriole and have to be removed from the mother centriole to allow ciliogenesis to proceed ^5,75^. We therefore determined the number of cells containing only one centrsomal dot of CP110 after 8 hours of serum withdrawal but found no difference between control- and BFA-treated RPE1 cells (Fig. 8d). This result suggests that the Golgi apparatus is not required for the earliest stage of ciliogenesis. Next, we analyzed whether EGFP-Rab8a was still recruited to the centrosome or cilium after incubation of RPE1 cells with BFA. As we have shown above, Rab8a apparently is required from the partial doughnut stage onwards. After 4 hours of serum starvation, Golgi disruption resulted in a less frequent accumulation of Rab8a at the centrosome or cilium (Fig. 8e) indicating that the Golgi apparatus is most likely important for more advanced stages of ciliogenesis to develop.

To more specifically determine the stage of ciliogenesis in which the Golgi apparatus is involved we subjected centrosomal regions of BFA-treated RPE1 cells stably expressing centrin1-EGFP to STEM tomography. At the same time, cells were incubated for 15 min with WGA-HRP to assess whether the accumulation of endocytosed membrane material at the mother centriole is perturbed upon disruption of the Golgi apparatus. Such a result would suggest that endocytosed membrane material traffics via the Golgi apparatus to the nascent ciliary membrane. This hypothesis was not confirmed by our data because we could not find any difference between BFA- and solvent-treated cells regarding the labeling of the nascent and mature ciliary membrane with WGA-HRP (Fig. 8f-j), which indicates that both the Golgi apparatus and endocytosed membrane material contribute to cilia formation independently of each other. Our argument is further supported by the fact that in solvent-treated cells we never observed Golgi stacks labeled with WGA-HRP. However, further analysis of the tomograms revealed a higher number of cells with earlier stages of cilia formation in BFA-treated cells compared to control-treated cells (Fig. 8k). Another surprising effect upon BFA treatment concerned the orientation of the mother centriole. While in solvent-treated cells the longitudinal axis of the mother centriole was preferentially oriented in a parallel fashion to the basal plasma membrane, it was oriented almost in a perpendicular fashion in BFA-treated cells with the distal end of the mother centriole pointing towards the basal plasma membrane (Fig. 8l).

## Discussion

### Discovery of novel intermediate stages of cilia formation

Based on our three-dimensional ultrastructural data we introduce a more refined model for the intracellular cilia assembly pathway that includes novel intermediate stages (Fig. 9). Beside the already described distal appendage vesicles, we also identified novel elongated membranous structures docked to distal appendages which we call distal appendage tubules. Both these structures are the first ultrastructural markers of cilia formation. They randomly dock to one or several distal appendages without any recognizable order. Fusion events between neighboring distal appendage vesicles (or tubules), and between distal appendage vesicles/tubules and incoming vesicles start before all distal appendages have been occupied. Our data support the lateral fusion of distal appendage vesicles and tubules to build a (partial) doughnut-shaped membrane structure (Fig. 1b and Fig. 9).

**Figure 9.**
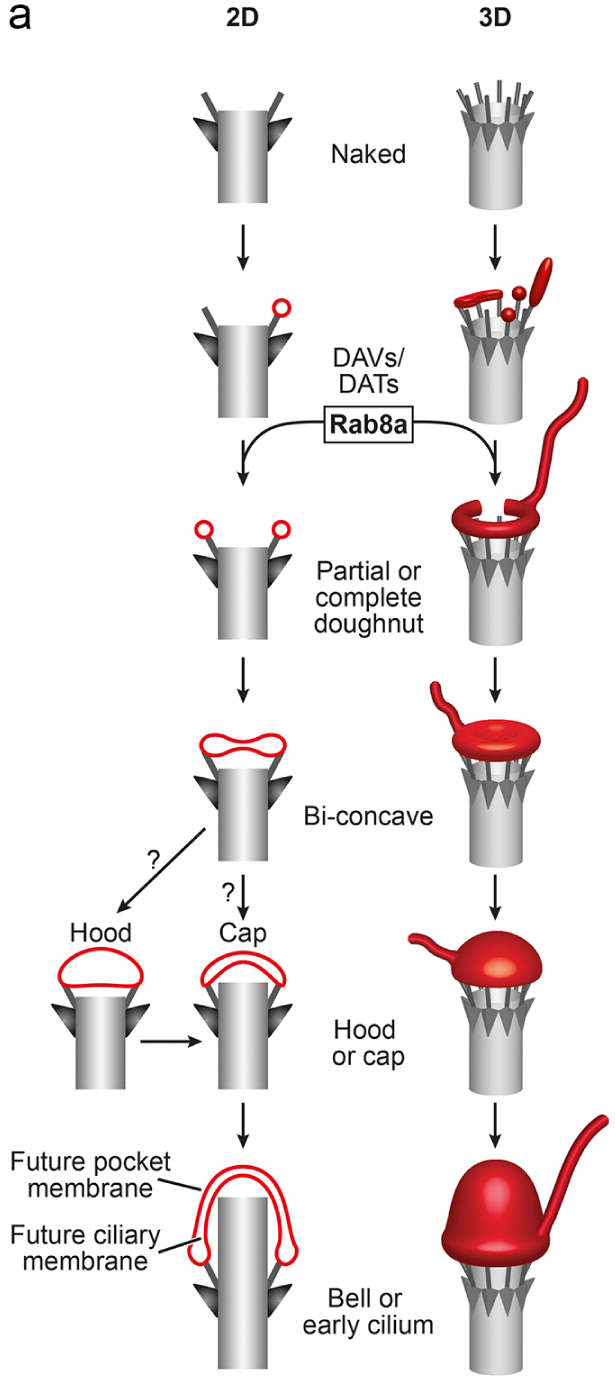
Novel model depicting the ultrastructural stages of the intracellular pathway of ciliogenesis. Vesicular and tubular structures associate with distal appendages of the mother centriole to form distal appendage vesicles and tubules (DAVs and DATs). These distal appendage vesicles and tubules fuse side by side, thereby developing into partial and complete doughnut-shaped membrane structures. Subsequently the cavity in the center is filled resulting in a bi-concave structure, out of which hood- and cap-like structures form. It is not clear whether ciliogenesis proceeds necessarily through the hood stage or whether the cap stage directly follows the bi-concave stage. From the partial doughnut stage onwards, tubular extensions of the nascent cilium can be observed. The mother centriole is drawn in grey and ciliary membrane structures in red. The left panel shows a longitudinal section and the right panel a corresponding three-dimensional view.

The rather frequent detection of the partial doughnut stage suggests that the formation and closure of a membranous ring is a time-consuming step. Comparable toroid membrane structures are pores in the nuclear envelope ^76^. Proteins mediating membrane bending, such as those containing a BAR (<u>B</u>in, <u>a</u>mphiphysin, <u>R</u>vs) domain ^77^ or an amphipathic helix ^78^, are apparently required for the formation and stabilization of membrane curvature. The F-BAR domain-containing proteins PACSIN1 and PACSIN2 (also known as syndapin 1 and 2) ^27^ and FAM92 ^79^ that were shown to be involved in cilia formation are promising candidates for the formation of doughnut-shaped membrane structures. Interestingly, we rarely detected complete doughnut-shaped membrane structures in any of our experimental conditions, which may result from the trivial reason that the whole membrane structure was not contained in the analyzed semi-thin sections. On the other hand, the complete doughnut-shaped membrane structure may represent a very transient stage and therefore is rarely detectable. It is remarkable that partial and complete ring-shaped membrane structures have not been identified during ciliogenesis so far. We believe that this is due to the limited information provided by the two-dimensional analysis of ultrathin sections. Depending on the plane of section a doughnut-shaped membrane structure was most likely misinterpreted as two distal appendage vesicles when the center of the doughnut was sectioned (e.g. Fig. 2b or Fig. 4h) or as a ciliary vesicle when the periphery of the doughnut was sectioned (e.g. Fig. 2b or Fig. 4h). It is noteworthy that we never identified a round ciliary vesicle in any cell.

Subsequent to the formation of a doughnut-shaped membrane structure, the hole in its center is filled in a centripetal fashion to build a bi-concave membrane structure. Further growth leads to a hood-shaped and then a cap-shaped membrane structure (Fig. 9). At this point of our analysis it has to be left open whether ciliogenesis necessarily proceeds through the hood stage to reach the cap stage or whether the cap stage can immediately follow the bi-concave stage. Finally, the axoneme starts to grow out forming a bell-shaped membrane structure and finally the cilium (Fig. 9). We would like to point out that not all nascent cilia strictly developed according to this model. Instead, we also observed membrane structures like an incomplete hood or cap not covering the complete distal end of the basal body, indicating that fusion events can also take place towards the center to fill the central hole before the ring is completely closed. This might account for the rare detection of the complete doughnut and the bi-concave stage as well.

The partial doughnut-shaped membrane structure is the first stage at which Rab8a localizes to the nascent primary cilium. Thus, Rab8a most likely functions earlier than during the extension of the ciliary vesicle as has been proposed based on depletion experiments ^5^. Considering the data presented in our current study, Rab8a may contribute to the closure of the doughnut-shaped membrane structure. EHD1 and EHD3, SNAP29 ^5^, PACSIN1 and PACSIN2 ^27^, and Rab34 ^80,81^ have been suggested to function during fusion of distal appendage vesicles while Cep97/CP110 is removed from the distal end of the mother centriole during this process ^75^. We speculate that EHD1, EHD3, SNAP29, PACSIN1, PACSIN2 and Rab34 may act during initial steps of ring formation as they were either shown to be recruited to the developing cilium earlier than Rab8a or are required for the association of Rab8a with the nascent cilium ^5,27,80,81^.

### Delivery and removal of membrane material

Depletion ^19^ and knock-out ^82^ of the distal appendage protein Cep164 ^83^ leads to an impaired docking of vesicles to distal appendages. However, it is not understood by what mechanism incoming membrane material attaches to these structures. Our three-dimensional structural analysis enabled us to detect filaments extending between the distal appendages and attached ciliary membrane structures. Similar filaments were observed between already attached ciliary membrane structures and additional incoming vesicles. These filamentous structures typically had the length of around 20-30 nm as has been described for tethers that bridge acceptor and donor membranes ^84,85^. Tethers that have been implicated in ciliogenesis are the homotypic fusion and vacuole protein sorting (HOPS)-tethering complex ^86,87^, the transport protein particle (TRAPP) II complex ^6,88^ and the exocyst complex ^89–91^.

As previously shown ^27^, we regularly observed tubules connected to the (future) ciliary pocket membrane. Their formation depends on EHD1 and the F-BAR proteins PACSIN1 and PACSIN2, and they were shown to connect the nascent cilium with the plasma membrane ^27^. In our study we detected such tubules from the partial doughnut stage onwards. Some of them ended in a blind fashion in the cytoplasm while others connected the ciliary membrane compartment to the plasma membrane and thus to the extracellular space. In many instances, however, we could not localize the distal ends of those tubules because they lay outside of the tomographed sections. The function of those tubules has to be left open just as the function of vesicles inside the lumen of centrioles which have been described many years ago ^2,29^. Despite our extensive and systematic analysis their role during ciliogenesis, if any, remains a puzzle.

On the outgoing side we identified clathrin-coated buds during very early stages of cilia formation, i.e. on distal appendage tubules and on partial doughnut-shaped membrane structures. This suggests that during ciliary growth membrane material is already removed to allow a continuous turnover of the developing ciliary membrane and not only at established primary cilia where clathrin-coated pits were described at the ciliary pocket membrane ^1,28^.

### Endocytotic processes mediated by GRAF1 contribute membrane material to the developing ciliary membrane compartment

A long-lasting question in the cilia field concerns the compartment(s) contributing to the establishment of the ciliary membrane and thus to ciliogenesis. Here, we provide evidence that endocytotic processes deliver membrane material to the growing ciliary membrane compartment. In principle, plasma membrane-derived material can reach the ciliary membrane compartment via a direct endocytosis route, by passing through the pericentriolar recycling compartment or through tubules connecting the plasma membrane with the nascent ciliary membrane ^27^. We consider the last possibility less likely because we identified several WGA-HRP-labeled early stages of ciliogenesis for which we were able to exclude a connection with the plasma membrane.

Our interference with any of the conventional endocytosis routes did not perturb ciliogenesis in RPE1 cells. This is in agreement with previous reports in which inhibition of clathrin-mediated endocytosis showed no negative effect on ciliogenesis ^1,92^. Also, our interference with the caveolae-dependent endocytosis pathway by pharmacological depletion of cholesterol or by overexpression of dominant-negative caveolin1 mutants did not impair ciliogenesis. Consistently, the overexpression of wild-type and mutant caveolin1 in RPE1 cells ^93^ and the silencing of caveolin1 in RPE1, IMCD3 and MDCK cells affected ciliary length without altering the percentage of ciliated cells ^94^. In contrast, in zebrafish and IMCD3 cells pharmacological depletion of cholesterol led to a decrease in cilia number and length ^95^, indicating species- and cell type-specific differences regarding the requirement of cholesterol and thus possibly caveolae for ciliogenesis.

In our study we identified the endocytosis-associated protein GRAF1 as a novel regulator of ciliogenesis. Its depletion led to an accumulation of cells in early ciliogenesis with the distal appendage vesicle/tubule stage being the most abundant. At the same time the delivery and/or recruitment of endocytosed membranes to the ciliary membrane compartment was markedly reduced. GRAF1 contains an NH_2_-terminal BAR domain, a PH (<u>p</u>leckstrin <u>h</u>omology) domain binding to lipids, an SH3 (<u>S</u>rc <u>h</u>omology 3) domain binding to dynamin, and a COOH-terminal Rho-GAP (<u>G</u>TPase <u>a</u>ctivating <u>p</u>rotein) domain directed towards Cdc42 and RhoA ^96^. It is known to function at two different sites and accordingly during two distinct processes: Firstly, GRAF1 acts at the plasma membrane and mediates the generation of tubular endocytotic membranes during CLIC/GEEC endocytosis ^64^. Secondly, GRAF1 functions during vesiculation of the tubular compartment of the recycling endosome to allow the return of receptors and lipids to the plasma membrane ^97^. During vesiculation of the tubular recycling endosome GRAF1 acts in a complex with MICAL-L1 and EHD1 ^97^. MICAL-L1 also participates in recruiting EHD1 and Rab8a to tubular membranes of the recycling endosome ^98^. Additionally, EHD1 links PACSINs to intracellular membranes ^99^. While PACSIN2 drives the tubulation of the recycling endosome ^100^, EHD1 and GRAF1 mediate the subsequent fission of vesicles from these tubules ^97^. Interestingly, these interaction partners of GRAF1 have all been linked to cilia formation ^5,6,27,101^. Moreover, the GTPase Rab11a also functions at the recycling endosome ^8–10^ and during ciliogenesis ^6,32,102^.

In principle, GRAF1 may act at the plasma membrane, the recycling endosome and/or the nascent cilium. On the one hand, GRAF1 could perform its function in ciliogenesis during CLIC formation at the plasma membrane and thus regulate the delivery of endocytic vesicles to the centrosomal region. On the other hand, GRAF1 could regulate the vesiculation of the tubular recycling endosome thereby generating vesicles destined not only for the plasma membrane but also for the ciliary membrane compartment. In this case, vesicles contributing to ciliogenesis would originate from the tubular recycling endosome. In a third scenario, GRAF1 could cooperate with EHDs and PACSINs at the developing ciliary membrane compartment itself and may stabilize the partial and complete doughnut-shaped membrane structure via its BAR domain. Indeed, ciliogenesis progression towards partial doughnut-shaped membrane structures was impaired when GRAF1 was depleted, suggesting that GRAF1 contributes to ring formation.

### The Golgi apparatus supports ciliogenesis independently of endocytotic processes

Inhibition of the Golgi apparatus by three small molecules resulted in less ciliated cells, which strongly supports the involvement of this organelle in the development of primary cilia. It is remarkable that various proteins such as the golgins giantin ^103,104^ and GMAP210 ^105^ as well as the intraflagellar transport component IFT20 ^106^ simultaneously play a role during Golgi function and cilia assembly. In our experiments, a collapse of the Golgi apparatus did not prevent the docking of distal appendage vesicles and tubules. In agreement with this finding, CP110 removal from the distal end of the mother centriole, which is necessary for ciliogenesis following the distal appendage vesicle stage, was not perturbed upon disruption of the Golgi apparatus but the centrosomal recruitment of Rab8a, that we have found to localize to the developing ciliary membrane compartment from the partial doughnut stage onwards, was inhibited.

The fact that WGA-HRP-labeled membranes still reached the developing cilium after disruption of the Golgi apparatus argues against the possibility that endocytosed membranes traffic retrogradely to the trans-Golgi network and reach the centrosomal region from there. Furthermore, this finding argues that the Golgi apparatus and endocytosis act independently of each other during ciliogenesis. At this stage it is not possible to decide whether the lower number of ciliated cells after disruption of the Golgi apparatus results from a reduced delivery of membrane material or from a misorientation of the mother centriole or both. Moreover, we do not know whether the mother centriole is not properly oriented because ciliogenesis does not progress or if proper orientation is required for ciliogenesis to proceed.

In summary, our study provides conclusive evidence that both GRAF1-dependent endocytotic processes and the Golgi apparatus independently support ciliogenesis.

## Materials and methods

### Plasmids

The cDNA for the GDP-locked mutant of human Rab5a [Rab5a (S34N)] was kindly provided by Stefan Linder (University Medical Center Eppendorf, Hamburg, Germany). The cDNA for the GDP-locked mutant of human Rab11a [Rab11a (S25N)] was kindly provided by Guiscard Seebohm (University Hospital Münster, Münster, Germany). The coding regions of the respective cDNAs were cloned into pEGFP-C1 (Clontech). An expression plasmid for the GDP-locked mutant of rat dynamin2 [dynamin2 (K44A) in pEGFP-N1] was kindly provided by Mark A. McNiven ^46^ (Mayo Clinic, Rochester, MN, USA). Expression plasmids for human Rab8a in pEGFP-C1 and pmCherry-C1 were a kind gift from Gislene Pereira ^19^ (Center for Organismal Studies, University of Heidelberg, Heidelberg, Germany). An expression plasmid encoding the COOH-terminal fragment of rat AP180 (isoform X2, aa 514-897) in pEGFP-C3 was a kind gift from Harvey T. McMahon (MRC Laboratory of Molecular Biology, Cambridge, UK) ^48^. The cDNA for human EPS15 was kindly provided by Alexandre Benmerah (INSERM, Paris, France) and was used to clone a truncated version lacking amino acids 1-528 [EPS15 (DIII)] into pEGFP-C1. The cDNA for human caveolin1 was kindly provided by Jeffrey E. Pessin (University of Iowa, Iowa City, Iowa, USA) and cloned into pEGFP-N1. The point mutation P132L was generated by site-directed mutagenesis. A truncated version of caveolin1 lacking amino acids 1-81 [caveolin1 (ΔN1-81)-EGFP] was generated using a PCR-based strategy again in pEGFP-N1. pMXs-Puro-ARHGAP26 containing the cDNA for human GRAF1 (splice version GRAF1b) was a gift from Axel Hillmer (Addgene plasmid # 69467; http://n2t.net/addgene:69467; RRID:Addgene_69467). The coding region of GRAF1 was cloned into pEGFP-C1. A mutant of GRAF1 resistant to RNA interference [GRAF1 (rescue)] was generated by site-directed mutagenesis and contains five silent point mutations. All constructs were confirmed by sequencing.

### Antibodies

The following primary antibodies were used for indirect immunofluorescence: rabbit polyclonal anti-Arl13b antibody (diluted 1:2,000; Proteintech, cat. no. 17711-1-AP), mouse monoclonal anti-γ-tubulin antibody GTU-88 (diluted 1:2,000; Sigma-Aldrich, cat. no. T6557), and mouse monoclonal anti-CP110 antibody 140-195-5 (diluted 1:500; Sigma-Aldrich, cat. no. MABT1354). The following primary antibodies were used for Western blot analysis: rabbit polyclonal anti-ARHGAP26 antibody (diluted 1:100; Sigma-Aldrich, cat. no. HPA035107), rabbit polyclonal anti-EGFP antibody (1:1,500; Invitrogen, cat. no. A-6455), and mouse monoclonal anti-actin antibody C4 (1:1,000; Sigma-Aldrich, cat. no. MAB1501). Secondary antibodies for immunofluorescence analysis were Alexa Fluor 350-coupled anti-mouse IgG (diluted 1:80; Invitrogen, cat. no. A11045), Alexa Fluor 488-coupled anti-rabbit IgG (diluted 1:600; Invitrogen, cat. no. A11034), Alexa Fluor 568-coupled anti-mouse IgG (diluted 1:600; Invitrogen, cat. no. A10037), Alexa Fluor 568-coupled anti-rabbit IgG (diluted 1:600; Invitrogen, cat. no. A10042). Secondary antibodies for Western blot analysis were horseradish peroxidase-coupled anti-mouse IgG (diluted 1:15,000; Sigma-Aldrich, cat. no. A3682) and horseradish peroxidase-coupled anti-rabbit IgG (diluted 1:70,000; Sigma-Aldrich, cat. no. A0545).

### Cell culture

Human telomerase reverse transcriptase (hTERT)-immortalized retinal pigment epithelial (RPE1) cells, stably transfected hTERT-RPE1 cells constitutively producing centrin1-EGFP ^21^ (a kind gift of Gislene Pereira, Center for Organismal Studies, Heidelberg, Germany) and other stably transfected hTERT-RPE1 cell lines generated in this study were cultured in DMEM/Ham’s F12 medium supplemented with 10% fetal calf serum, 2 mM L-glutamine and 0.348% sodium bicarbonate (RPE1 cells for short from now on). All cell lines were grown at 37°C under humidified conditions and 5% CO_2_. RPE1 cells were transiently transfected with plasmid DNA using Fugene 6 (Promega) according to the manufacturer’s protocol and serum-starved 24 hours later for up to 24 hours before being fixed. To generate stably transfected RPE1 (centrin1-EGFP) cells additionally producing mCherry-Rab8a transfection was performed together with the plasmid pWE3 (ATCC plasmid # 87673) ^107^ carrying the puromycin resistance gene to allow subsequent selection with 2 µg/ml of puromycin for approximately 2 weeks. After FACS sorting by the BD FACSAria™ IIu cell sorting system, single clones were obtained by limiting dilution and analyzed by fluorescence microscopy. To generate a stably transfected RPE1 cell line constitutively producing EGFP-Rab8a, cells were transfected and selected with 800 µg/ml of G418 for approximately 2 weeks. Single clones were picked and analyzed by fluorescence microscopy. All cell lines were confirmed to be free of mycoplasma on a regular basis.

### Chemicals and inhibition assays

To inhibit endocytosis, endosomal trafficking, and to disrupt or inhibit the Golgi apparatus RPE1 cells were serum-deprived to induce cilia formation 48 h after plating. Simultaneously, the following compounds were added to the medium lacking FCS: 40 µM dynasore (Sigma-Aldrich, cat. no. D7693) or 0.08% DMSO, 0.5 mM methyl-β-cyclodextrin (Sigma-Aldrich, cat. no. C4555) or 5% medium without FCS, 50 nM bafilomycin A1 or 0.03125% DMSO (Sigma-Aldrich, cat. no. SML1661), 3 µM wortmannin (Sigma-Aldrich, cat. no. W3144) or 0.03% DMSO, 10 µM LY294002 (Selleckchem, cat. no. S1105) or 0.1% DMSO, 0.5 µg/ml of brefeldin A (Sigma-Aldrich, cat. no. B6542) or 0.005% methanol, 1 µM 30N12 (Chembridge, cat. no. 5175331) or 0.01% DMSO, 2 µM monensin (Sigma-Aldrich, cat. no. M5273) or 0.02% ethanol. Cells were fixed 0, 4, 8, 12, or 24 h after serum starvation and incubation with the respective compound or solvent.

### RNA interference

RPE1 and RPE1 (centrin1-EGFP) cells were transfected with 10 nM (final concentration) siRNA oligonucleotides using Lipofectamine RNAiMAx (Invitrogen) according to the manufacturer’s instructions. Cells were lysed or serum-starved 48 h after transfection and fixed 0, 12 or 24 h after serum deprivation. Silencer Select siRNAs were purchased from Invitrogen and targeted the following sequences: human Cep164 siRNA, 5’CAGGTGACATTTACTATTTCA-3’ ^83^, and human Graf1 siRNA, 5’- CTCATGATGTACCAGTTTCAA-3‘ (modified from ^64^). As a negative control, non-targeting Silencer Select Negative Control No. 1 siRNA was used (Invitrogen).

For rescue experiments RPE1 cells were transiently transfected with the plasmid pEGFP-C1/GRAF1 (rescue). The next day cells were transfected with the respective siRNAs. After 24 h cells were serum-deprived for another 24 h before being fixed or lysed.

### Cell lysis and Western blot

RPE1 cells were resuspended in RIPA buffer [50 mM Tris-HCl pH 8.0, 150 mM NaCl, 1% IGEPAL, 1% sodium deoxycholate, 0.1% SDS, 1 mM EDTA, 1 mM DTT, 2 mM Na_3_VO_4_, 1 mM PMSF, complete EDTA-free protease inhibitor cocktail (Roche)] and incubated on a rotating wheel for 20 min at 4°C before being centrifuged for 15 min at 21,500 *g*. The supernatant was recovered and the protein concentration determined with Bradford reagent (Roti-Quant, Roth). Aliquots of the lysates were boiled in SDS sample buffer and separated by SDS-PAGE. After transfer to a PVDF membrane, proteins of interest were detected with the respective antibodies and visualized by chemiluminescence (WesternBright Chemilumineszenz Substrat Quantum, Biozym Scientific) using a Fusion-FX7 imaging system (Vilber Lourmat).

### Indirect immunofluorescence

Cells were grown on 10 mm coverslips, fixed for 20 min in 1x PBS, 4% paraformaldehyde, quenched for 10 min in 1x PBS, 50 mM NH_4_Cl and permeabilized for 5 min in 1x PBS, 0.1% Triton X-100. After blocking for 30 min with 1x PBS, 0.5% BSA, 0.1% Triton X-100, cells were incubated for 1 h at 37°C with the primary antibody. Specific staining was detected by incubating for 30 min at room temperature with Alexa Fluor–conjugated secondary antibodies. DNA was stained with 0.5 µg/ml of Hoechst 33258 in 1x PBS. Only 1.7 ng/ml of Hoechst 33258 was used when Alexa 350-secondary antibodies were applied. Coverslips were mounted on glass slides in Mowiol 4-88. Images were acquired as *z* stacks with 0.4 µm steps using an inverted microscope (Axiovert 200M, Carl Zeiss) equipped with a 40x Fluar NA 1.3 oil immersion objective (Carl Zeiss), a 63x Plan Apochromat NA 1.4 oil immersion objective (Carl Zeiss), a sCMOS pco.edge camera (PCO AG) and VisiView software (Visitron Sysmtes GmbH). At least four fields per condition were imaged from two different coverslips. Analysis of fluorescence images and quantification of cilia was manually conducted with Fiji/ImageJ^108,109^.

### Correlative light and electron microscopy (CLEM)

RPE1 (centrin1-EGFP, mCherry-Rab8a) cells and RPE1 (centrin1-EGFP) cells transiently expressing mCherry were grown at a low density on gridded glass bottom dishes with an alphanumerical code (MatTek) to allow for the subsequent identificaton of specific cells in Epon blocks and sections. Cells were fixed for 30 min at room temperature with 4% paraformaldehyde in 0.1 M sodium cacodylate pH 7.4 and kept in 0.1 M sodium cacodylate pH 7.4 during light microscopy. Regions with optimal cell density were documented with an Axiovert 200M microscope (Carl Zeiss) equipped with a 40x Fluar NA 1.3 oil immersion objective (Carl Zeiss). Seventeen planes spanning a total of 6.4 µm in the *z* axis were typically recorded. Brightfield images were recorded with a 16x Plan-Neofluar NA 0.5 oil immersion objective (Carl Zeiss). After light microscopy, cells were additionally fixed for 5 min at room temperature with 2% glutardialdehyde in 0.1 M sodium cacodylate pH 7.4 and another 55 min or overnight on ice. The cells were contrasted for 30 min at 4°C with 1% OsO_4_ in 0.1 M sodium cacodylate pH 7.4 and overnight at 4°C with a 1% aqueous uranyl acetate solution. Finally cells were embedded in Epon. Fluorescence images were analyzed with Fiji/ImageJ ^108,109^ to identify centrosomal regions. After selected regions of interest were relocated in the Epon block based on the pattern from the dish’s gridded bottom, 600 nm thick serial sections were prepared in parallel to the basal surface of the cell layer with an ultramicrotome EM UC6 (Leica Microsystems) and collected on pioloform-coated slot copper grids (Plano).

### WGA-HRP uptake and CLEM

RPE1 (centrin1-EGFP) cells were grown at a low density on gridded glass bottom dishes (MatTek). For depletion experiments, cells were transfected with siRNA during seeding as described above and serum-starved 48 h later for 4 or 12 h. When the Golgi apparatus was disrupted, cells were treated with 0.005% methanol or 0.5 µg/ml of BFA for 2 h in medium with serum before they were serum-deprived for 4 h. For each condition, cells were incubated for the last 15 min of serum starvation with 30 µg/ml of WGA-HRP (Vector Labs, cat. no. PL-1026-2) at 37°C. Cells were fixed for 30 min at room temperature in 4% paraformaldehyde, 0.1 M sodium cacodylate pH 7.4. To visualize nuclei, cells were stained for 8 min at room temperature with 170 ng/ml of Hoechst 33342 in 0.1 M sodium cacodylate pH 7.4. Light microscopy was carried out in 0.1 M sodium cacodylate pH 7.4 as described above. Subsequently cells were fixed for 5 min at room temperature and then overnight at 4°C in 2% glutardialdehyde, 0.1 M sodium cacodylate pH 7.4. To visualize peroxidase activity, cells were incubated at constant agitation for 15 min on ice with 0.5 mg/ml of diaminobenzidine in 50 mM Tris-HCl pH 7.6 in the dark. Then H_2_O_2_ was added to a final concentration of 0.02% and cells were incubated another 30 min. Finally, cells were contrasted for 30 min at 4°C with 1% OsO_4_, 0.1 M sodium cacodylate pH 7.4 and for 1 h at 4°C with 1% aqueous uranyl acetate, dehydrated in a graded ethanol series and embedded in Epon. Fluorescence images were analyzed and centrosomal regions with not more than two centrin1-EGFP-positive centrioles were selected for further three-dimensional investigation. Samples in resins were processed and 900 nm sections prepared as mentioned earlier.

### Scanning transmission electron microscopy (STEM) and electron tomography

Cells previously documented by fluorescence microscopy were re-located by transmission electron microscopy using a JEM-2100F field emission electron microscope operated at 200 kV (JEOL). Micrographs os different magnifications were recorded with a TemCamF416 camera (TVIPS) that was operated with the software EM-MENU 5 (TVIPS). Correlation of transmission electron micrographs with brightfield light microscopy and fluorescence images using the TVIPS CLEM software facilitated the re-identification process of centrosomal regions in thick Epon sections. To increase the electrical conductivity of specimens during STEM tomography, a carbon layer of ∼3 nm thickness was applied to the Epon sections by a high vacuum carbon coater (208 Carbon Turbo; Cressington Scientific Instruments). Colloidal protein A-gold particles with a diameter of 15 nm (purchased from George Posthuma, University of Utrecht, Netherlands) were applied to both sides of the grids to serve as fiducial markers for alignment during the reconstruction process ^22,23^. After plasma cleaning (PDC-3XG Harrick Plasma), tilt series were recorded by STEM tomography ^23^ with a nominal magnification of 80,000x, 120,000x and 200,000x. The pixel size was 6.69, 2.23, and 1.34 nm, respectively, and the fields of view were approximately 6.85, 4.6 or 2.7 µm in the *x* and *y* directions, respectively. A series of 90 images from +66° to -66° (non-linear increment, Saxton scheme) ^23^ was recorded using a STEM bright-field detector (JEOL), a Universal Scan Generator (TVIPS) and the software EM-TOOLS (TVIPS). Dual-axis STEM tomograms with a nominal magnification of 200,000x were additionally recorded for selected sample areas ^23^. Three-dimensional reconstructions were calculated with IMOD (4.12) ^110^ using fiducial markers for alignment. Tomograms were generated by weighted back projection including a SIRT (simultaneous iterative reconstruction technique)-like filter equivalent to 15 iterations.

### Re-identification of centrosomal regions of interest from fluorescence imaging in electron microscopic micrographs

Cells previously imaged by fluorescence microscopy were re-identified in the electron microscope by using the alphanumeric grid pattern, the outlines of the cell, the shape and position of the nuclei, and the heterochromatin structures. To ensure that the mCherry-Rab8a signal was indeed located close to the distal end of the mother centriole, tomograms were appropriately rotated to align brightfield, fluorescence and transmission electron micrographs. For better visualization, overlays between fluorescence images and STEM tomograms were generated by eC-CLEM ^25^, a software plugin of the Icy platform ^111^, with the two centrioles serving as landmarks. For each overlay between fluorescence image and STEM tomogram the correlation accuracy was calculated ^22^.

### Tomogram segmentation to generate 3D models of nascent primary cilia

Raw tomographic data were binned 2x2 using IMOD ^110^ and slices outside the sample volume were removed. The following structures of the nascent primary cilia were highlighted by manual segmentation using the AMIRA software package (Visage Imaging): centriolar and axonemal microtubules, intracentriolar vesicles, subdistal appendages, distal appendages, membrane structures docked to distal appendages, filamentous connections between distal appendages and docked membrane structures, membrane structures touching other membrane structures, membrane structures connected to other membrane structures via filaments, filamentous connections between membrane structures, and the plasma membrane. Smoothing was not applied.

### Angle measurements

Angles between the longitudinal axis of the mother centriole and the sectional plane were measured using the AMIRA software package (Visage Imaging). Only tomograms that contained complete mother centrioles were included. Raw data of reconstructed tomograms with a nominal magnification of 80,000x or 200,000x were binned 2x2 with IMOD ^110^. Two anchoring points in the center of the mother centriole were positioned at its distal and its proximal end and connected by interpolation. The correct position of this line in the center of the mother centriole was verified in all three views (*xy, xz, yz*) and accordingly adjusted if required. The angle between this line and the sectional plane was measured with the angle measurement tool in different views after which the lowest value was considered. Angles of mother centrioles whose distal end was directed towards the apical plasma membrane were given a positive sign, while angles of mother centrioles whose distal end pointed towards the basal plasma membrane received a negative sign.

### Size measurements of intracentriolar vesicles

The size of intracentriolar vesicles was measured using the IMOD software package after 2x2 binning of the tomograms ^110^. Only tomograms containing the entire centriole were included. The slice of the tomogram showing the respective vesicle in its largest dimension was used for measurement, and the diameter of each vesicle was measured in the *x* and *y* dimensions.

### Statistical analysis

When applicable, results are presented as means + standard deviation of the number of independent experiments. Horizontal lines of box-and-whisker plots show the 25^th^, 50^th^ (median) and 75^th^ percentiles, whiskers extend to minimum and maximum values; the mean value and outliers are indicated. Statistical analysis was performed using SPSS. To test for normal distribution, the Shapiro-Wilk test was applied. When two normally distributed data sets were compared, a two-tailed unpaired student’s *t*-test was used. In the case of two data sets which were not normally distributed, a Mann-Whitney U-test was employed. When more than two normally distributed datasets were compared, one-way ANOVA followed by Dunnett’s or Tukey’s post hoc test was used. The Kruskal-Wallis test followed by Dunn’s test was applied for more than two data sets that were not normally distributed. Pearson’s Chi-square test was applied when the distribution of a categorical variable was compared between two samples.

## Acknowledgements

We thank Alexandre Benmerah, Axel Hillmer, Stefan Linder, Harvey T. McMahon, Mark A. McNiven, Gislene Pereira, Jeffrey E. Pessin, and Guiscard Seebohm for sharing reagents. We are very grateful for the reconstruction of tomograms and the arrangement of figures by Ton Maurer. We thank the student assistants Selina Kalesse, Selin Kuecuekoktay and Alexandra Schuldt for excellent technical support. The expertise of the members of the FACS Core Facility at the Leibniz Institute for Immunotherapy was instrumental for the isolation of transfected cells.

This work was supported by the Deutsche Forschungsgemeinschaft (DFG, German Research Foundation), project number 387509280 (SFB 1350) and project number 509149993 (TRR 374).

## References

1 Molla-Herman, A. et al. The ciliary pocket: An endocytic membrane domain at the base of primary and motile cilia. J. Cell Sci. 123, 1785–1795 (2010).

2 Sorokin, S. Centrioles and the formation of rudimentary cilia by fibroblasts and smooth muscle cells. J. Cell Biol. 15, 363–377 (1962).

3 Dingle, A. D. & Fulton, C. Development of the flagellar apparatus of *Naegleria*. J. Cell Biol. 31, 43–54 (1966).

4 Archer, F. L. & Wheatley, D. N. Cilia in cell-cultured fibroblasts. II. Incidence in mitotic and post-mitotic BHK 21/C13 fibroblasts. J. Anat. 109, 277–292 (1971).

5 Lu, Q. et al. Early steps in primary cilium assembly require EHD1/EHD3-dependent ciliary vesicle formation. Nat. Cell Biol. 17, 228–240 (2015).

6 Westlake, C. J. et al. Primary cilia membrane assembly is initiated by Rab11 and transport protein particle II (TRAPPII) complex-dependent trafficking of Rabin8 to the centrosome. Proc. Natl. Acad. Sci. USA 108, 2759–2764 (2011).

7 Oguchi, M. E., Okuyama, K., Homma, Y. & Fukuda, M. A comprehensive analysis of Rab GTPases reveals a role for Rab34 in serum starvation-induced primary ciliogenesis. J. Biol. Chem. 295, 12674–12685 (2020).

8 Huotari, J. & Helenius, A. Endosome maturation. EMBO J. 30, 3481–3500 (2011).

9 Stenmark, H. Rab GTPases as coordinators of vesicle traffic. Nat. Rev. Mol. Cell Biol. 0, 513–525 (2009).

10 Wandinger-Ness, A. & Zerial, M. Rab proteins and the compartmentalization of the endosomal system. Cold Spring Harb. Perspect. Biol. 6, a022616 (2014).

11 Kim, J. et al. Functional genomic screen for modulators of ciliogenesis and cilium length. Nature 464, 1048–1051 (2010).

12 Inoue, H., Ha, V. L., Prekeris, R. & Randazzo, P. A. Arf GTPase-activating protein ASAP1 interacts with Rab11 effector FIP3 and regulates pericentrosomal localization of transferrin receptor-positive recycling endosome. Mol. Biol. Cell 19, 4224–4237 (2008).

13 Doyotte, A., Mironov, A., McKenzie, E. & Woodman, P. The Bro1-related protein HD-PTP/PTPN23 is required for endosomal cargo sorting and multivesicular body morphogenesis. Proc. Natl. Acad. Sci. USA 105, 6308–6313 (2008).

14 Grant, B. D. & Caplan, S. Mechanisms of EHD/RME-1 protein function in endocytic transport. Traffic 9, 2043–2052 (2008).

15 Hsiao, Y.-C. et al. Ahi1, whose human ortholog is mutated in Joubert syndrome, is required for Rab8a localization, ciliogenesis and vesicle trafficking. Hum. Mol. Genet. 18, 3926–3941 (2009).

16 Troilo, A. et al. HIF1α deubiquitination by USP8 is essential for ciliogenesis in normoxia. EMBO Rep. 15, 77–85 (2014).

17 Stenmark, H. et al. Inhibition of rab5 GTPase activity stimulates membrane fusion in endocytosis. EMBO J. 13, 1287–1296 (1994).

18 Kumar, D. et al. A ciliopathy complex builds distal appendages to initiate ciliogenesis. J. Cell Biol. 220, e202011133 (2021).

19 Schmidt, K. N. et al. Cep164 mediates vesicular docking to the mother centriole during early steps of ciliogenesis. J. Cell Biol. 199, 1083–1101 (2012).

20 Wu, C.-T., Chen, H.-Y. & Tang, T. K. Myosin-Va is required for preciliary vesicle transportation to the mother centriole during ciliogenesis. Nat. Cell Biol. 20, 175–185 (2018).

21 Uetake, Y. et al. Cell cycle progression and de novo centriole assembly after centrosomal removal in untransformed human cells. J. Cell Biol. 176, 173–182 (2007).

22 Buerger, K. et al. On-section correlative light and electron microscopy of large cellular volumes using STEM tomography. Methods Cell Biol. 162, 171–203 (2021).

23 Rachel, R. et al. Dual-axis STEM tomography at 200 kV: Setup, performance, limitations. J. Struct. Biol. 211, 107551 (2020).

24 Kremer, J. R., Mastronarde, D. N. & McIntosh, J. R. Computer visualization of three-dimensional image data using IMOD. J. Struct. Biol. 116, 71–76 (1996).

25 Paul-Gilloteaux, P. et al. eC-CLEM: Flexible multidimensional registration software for correlative microscopies. Nat. Methods 14, 102–103 (2017).

26 Witzgall, R. Golgi bypass of ciliary proteins. Sem. Cell Dev. Biol. (2018).

27 Insinna, C. et al. Investigation of F-BAR domain PACSIN proteins uncovers membrane tubulation function in cilia assembly and transport. Nat. Commun. 10, 428 (2019).

28 Rattner, J. B., Sciore, P., Ou, Y., van der Hoorn, F. A. & Lo, I. K. Y. Primary cilia in fibroblast-like type B synoviocytes lie within a cilium pit: A site of endocytosis. Histol. Histopathol. 25, 865–875 (2010).

29 Poole, A. C., Flint, M. H. & Beaumont, B. W. Analysis of the morphology and function of primary cilia in connective tissues: A cellular cybernetic probe? Cell Motil. 5, 175–193 (1985).

30 Nachury, M. V. et al. A core complex of BBS proteins cooperates with the GTPase Rab8 to promote ciliary membrane biogenesis. Cell 129, 1201–1213 (2007).

31 Follit, J. A., Li, L., Vucica, Y. & Pazour, G. J. The cytoplasmic tail of fibrocystin contains a ciliary targeting sequence. J. Cell Biol. 188, 21–28 (2010).

32 Knödler, A. et al. Coordination of Rab8 and Rab11 in primary ciliogenesis. Proc. Natl. Acad. Sci. USA 107, 6346–6351 (2010).

33 Ward, H. H. et al. A conserved signal and GTPase complex are required for the ciliary transport of polycystin-1. Mol. Biol. Cell 22, 3289–3305 (2011).

34 Bigge, B. M., Rosenthal, N. E. & Avasthi, P. Initial ciliary assembly in *Chlamydomonas* requires Arp2/3 complex-dependent endocytosis. Mol. Biol. Cell 34, ar24 (2023).

35 Nagata, Y. & Burger, M. M. Wheat germ agglutinin. Molecular characteristics and specificity for sugar binding. J. Biol. Chem. 249, 3116–3122 (1974).

36 Privat, J.-P., Delmotte, F. & Monsigny, M. Protein-sugar interactions. Association of β-(1 leads to 4) linked *N*-acetyl-D-glucosamine oligomer derivatives with wheat germ agglutinin (lectin). FEBS Lett. 46, 224–228 (1974).

37 Burger, M. M. & Goldberg, A. R. Identification of a tumor-specific determinant on neoplastic cell surfaces. Proc. Natl. Acad. Sci. USA 57, 359–366 (1967).

38 Greenaway, P. J. & LeVine, D. Binding of N-acetyl-neuraminic acid by wheat-germ agglutinin. Nat. New Biol. 241, 191–192 (1973).

39 Gonatas, N. K., Kim, S. U., Stieber, A. & Avrameas, S. Internalization of lectins in neuronal GERL. J. Cell Biol. 73, 1–13 (1977).

40 Pavelka, M., Neumüller, J. & Ellinger, A. Retrograde traffic in the biosynthetic-secretory route. Histochem. Cell Biol. 129, 277–288 (2008).

41 Vetterlein, M., Ellinger, A., Neumüller, J. & Pavelka, M. Golgi apparatus and TGN during endocytosis. Histochem. Cell Biol. 117, 143–150 (2002).

42 Kaksonen, M. & Roux, A. Mechanisms of clathrin-mediated endocytosis. Nat. Rev. Mol. Cell Biol. 19, 313–326 (2018).

43 Parton, R. G. Caveolae: Structure, function, and relationship to disease. Annu. Rev. Cell Dev. Biol. 34, 111–136 (2018).

44 Antonny, B. et al. Membrane fission by dynamin: What we know and what we need to know. EMBO J. 35, 2270–2284 (2016).

45 Macia, E. et al. Dynasore, a cell-permeable inhibitor of dynamin. Dev. Cell 10, 839–850 (2006).

46 Cao, H., Thompson, H. M., Krueger, E. W. & McNiven, M. A. Disruption of Golgi structure and function in mammalian cells expressing a mutant dynamin. J. Cell Sci. 113, 1993–2002 (2000).

47 Benmerah, A. et al. AP-2/Eps15 interaction is required for receptor-mediated endocytosis. J. Cell Biol. 140, 1055–1062 (1998).

48 Ford, M. G. J. et al. Simultaneous binding of PtdIns(4,5)P2 and clathrin by AP180 in the nucleation of clathrin lattices on membranes. Science 291, 1051–1055 (2001).

49 Klein, U., Gimpl, G. & Fahrenholz, F. Alteration of the myometrial plasma membrane cholesterol content with β-cyclodextrin modulates the binding affinity of the oxytocin receptor. Biochemistry 34, 13784–13793 (1995).

50 Mahammad, S. & Parmryd, I. Cholesterol depletion using methyl-β-cyclodextrin. Methods Mol. Biol. 1232, 91–102 (2015).

51 Pol, A. et al. A caveolin dominant negative mutant associates with lipid bodies and induces intracellular cholesterol imbalance. J. Cell Biol. 152, 1057–1070 (2001).

52 Lee, H. et al. Caveolin-1 mutations (P132L and null) and the pathogenesis of breast cancer. Caveolin-1 (P132L) behaves in a dominant-negative manner and caveolin-1 (-/-) null mice show mammary epithelial cell hyperplasia. Am. J. Pathol. 161, 1357–1369 (2002).

53 Bowman, E. J., Siebers, A. & Altendorf, K. Bafilomycins: A class of inhibitors of membrane ATPases from microorganisms, animal cells, and plant cells. Proc. Natl. Acad. Sci. USA 85, 7972–7976 (1988).

54 Raiborg, C., Schink, K. O. & Stenmark, H. Class III phosphatidylinositol 3-kinase and its catalytic product PtdIns3P in regulation of endocytic membrane traffic. FEBS J. 280, 2730–2742 (2013).

55 Okada, T., Sakuma, L., Fukui, Y., Hazeki, O. & Ui, M. Blockage of chemotactic peptide-induced stimulation of neutrophils by wortmannin as a result of selective inhibition of phosphatidylinositol 3-kinase. J. Biol. Chem. 269, 3563–3567 (1994).

56 Thelen, M., Wymann, M. P. & Langen, H. Wortmannin binds specifically to 1-phosphatidylinositol 3-kinase while inhibiting guanine nucleotide-binding protein-coupled receptor signaling in neutrophil leukocytes. Proc. Natl. Acad. Sci. USA 91, 4960–4964 (1994).

57 Gharbi, S. I. et al. Exploring the specificity of the PI3K family inhibitor LY294002. Biochem. J. 404, 15–21 (2007).

58 Vlahos, C. J., Matter, W. F., Hui, K. Y. & Brown, R. F. A specific inhibitor of phosphatidylinositol 3-kinase, 2-(4-morpholinyl)-8-phenyl-4H-1-benzopyran-4-one (LY294002). J. Biol. Chem. 269, 5241–5248 (1994).

59 Renard, H.-F. & Boucrot, E. Unconventional endocytic mechanisms. Curr. Opin. Cell Biol. 71, 120–129 (2021).

60 Thottacherry, J. J., Sathe, M., Prabhakara, C. & Mayor, S. Spoiled for choice: Diverse endocytic pathways function at the cell surface. Annu. Rev. Cell Dev. Biol. 35, 55–84 (2019).

61 Lakshminarayan, R. et al. Galectin-3 drives glycosphingolipid-dependent biogenesis of clathrin-independent carriers. Nat. Cell Biol. 16, 595–603 (2014).

62 Sabharanjak, S., Sharma, P., Parton, R. G. & Mayor, S. GPI-anchored proteins are delivered to recycling endosomes via a distinct cdc42-regulated, clathrin-independent pinocytic pathway. Dev. Cell 2, 411–423 (2002).

63 Francis, M. K. et al. Endocytic membrane turnover at the leading edge is driven by a transient interaction between Cdc42 and GRAF1. J. Cell Sci. 128, 4183–4195 (2015).

64 Lundmark, R. et al. The GTPase-activating protein GRAF1 regulates the CLIC/GEEC endocytic pathway. Curr. Biol. 18, 1802–1808 (2008).

65 Barr, F. A. Membrane traffic: Golgi stumbles over cilia. Curr. Biol. 19, R253–R255 (2009).

66 Masson, J. & El Ghouzzi, V. Golgi dysfunction in ciliopathies. Cells 11, 2773 (2022).

67 Stevenson, N. L. The factory, the antenna and the scaffold: The three-way interplay between the Golgi, cilium and extracellular matrix underlying tissue function. Biol. Open 12, bio059719 (2023).

68 Wheatley, D. N. Cilia and centrioles of the rat adrenal cortex. J. Anat. 101, 223–237 (1967).

69 Donaldson, J. G. & Jackson, C. L. Regulators and effectors of the ARF GTPases. Curr. Opin. Cell Biol. 12, 475–482 (2000).

70 Jackson, C. L. & Casanova, J. E. Turning on ARF: The Sec7 family of guanine-nucleotide-exchange factors. Trends Cell Biol. 10, 60–67 (2000).

71 Lippincott-Schwartz, J., Yuan, L. C., Bonifacino, J. S. & Klausner, R. D. Rapid redistribution of Golgi proteins into the ER in cells treated with brefeldin A: Evidence for membrane cycling from Golgi to ER. Cell 56, 801–813 (1989).

72 Ulmer, J. B. & Palade, G. E. Targeting and processing of glycophorins in murine erythroleukemia cells: Use of brefeldin A as a perturbant of intracellular traffic. Proc. Natl. Acad. Sci. USA 86, 6992–6996 (1989).

73 Mollenhauer, H. H., Morré, D. J. & Rowe, L. D. Alteration of intracellular traffic by monensin; mechanism, specificity and relationship to toxicity. Biochim. Biophys. Acta 1031, 225–246 (1990).

74 Yarrow, J. C., Feng, Y., Perlman, Z. E., Kirchhausen, T. & Mitchison, T. J. Phenotypic screening of small molecule libraries by high throughput cell imaging. Comb. Chem. High Throughput Screen. 6, 279–286 (2003).

75 Spektor, A., Tsang, W. Y., Khoo, D. & Dynlacht, B. D. Cep97 and CP110 suppress a cilia assembly program. Cell 130, 678–690 (2007).

76 Torbati, M., Lele, T. P. & Agrawal, A. Ultradonut topology of the nuclear envelope. Proc. Natl. Acad. Sci. USA 113, 11094–11099 (2016).

77 Simunovic, M., Voth, G. A., Callan-Jones, A. & Bassereau, P. When physics takes over: BAR proteins and membrane curvature. Trends Cell Biol. 25, 780–792 (2015).

78 Drin, G. & Antonny, B. Amphipathic helices and membrane curvature. FEBS Lett. 584, 1840–1847 (2010).

79 Li, F.-Q. et al. BAR domain-containing FAM92 proteins interact with Chibby1 to facilitate ciliogenesis. Mol. Cell. Biol. 36, 2668–2680 (2016).

80 Ganga, A. K. et al. Rab34 GTPase mediates ciliary membrane formation in the intracellular ciliogenesis pathway. Curr. Biol. 31, 2895–2905 (2021).

81 Stuck, M. W., Chong, W. M., Liao, J.-C. & Pazour, G. J. Rab34 is necessary for early stages of intracellular ciliogenesis. Curr. Biol. 31, 2887–2894 (2021).

82 Daly, O. M. et al. CEP164-null cells generated by genome editing show a ciliation defect with intact DNA repair capacity. J. Cell Sci. 129, 1769–1774 (2016).

83 Graser, S. et al. Cep164, a novel centriole appendage protein required for primary cilium formation. J. Cell Biol. 179, 321–330 (2007).

84 Szentgyörgyi, V. & Spang, A. Membrane tethers at a glance. J. Cell Sci. 136, jcs260471 (2023).

85 Ungermann, C. & Kümmel, D. Structure of membrane tethers and their role in fusion. Traffic 20, 479–490 (2019).

86 Iaconis, D. et al. The HOPS complex subunit VPS39 controls ciliogenesis through autophagy. Hum. Mol. Genet. 29, 1018–1029 (2020).

87 Jewett, C. E. et al. RAB19 directs cortical remodeling and membrane growth for primary ciliogenesis. Dev. Cell 56, 325–340 (2021).

88 Cuenca, A. et al. The C7orf43/TRAPPC14 component links the TRAPPII complex to Rabin8 for preciliary vesicle tethering at the mother centriole during ciliogenesis. J. Biol. Chem. 294, 15418–15434 (2019).

89 Chiba, S., Amagai, Y., Homma, Y., Fukuda, M. & Mizuno, K. NDR2-mediated Rabin8 phosphorylation is crucial for ciliogenesis by switching binding specificity from phosphatidylserine to Sec15. EMBO J. 32, 874–885 (2013).

90 Zuo, X., Fogelgren, B. & Lipschutz, J. H. The small GTPase Cdc42 is necessary for primary ciliogenesis in renal tubular epithelial cells. J. Biol. Chem. 286, 22469–22477 (2011).

91 Zuo, X., Guo, W. & Lipschutz, J. H. The exocyst protein Sec10 is necessary for primary ciliogenesis and cystogenesis in vitro. Mol. Biol. Cell 20, 2522–2529 (2009).

92 Saito, M. et al. Tctex-1 controls ciliary reabsorption by regulating branched actin polymerization and endocytosis. EMBO Rep. 18, 1460–1472 (2017).

93 Aslanyan, M. G. et al. A targeted multi-proteomics approach generates a blueprint of the ciliary ubiquitinome. Front. Cell Dev. Biol. 11, 1113656 (2023).

94 Rangel, L. et al. Caveolin-1α regulates primary cilium length by controlling RhoA GTPase activity. Sci. Rep. 9, 1116 (2019).

95 Maerz, L. D. et al. Pharmacological cholesterol depletion disturbs ciliogenesis and ciliary function in developing zebrafish. Commun. Biol. 2, 31 (2019).

96 Doherty, G. J. & Lundmark, R. GRAF1-dependent endocytosis. Biochem. Soc. Trans. 37, 1061–1065 (2009).

97 Cai, B., Xie, S., Caplan, S. & Naslavsky, N. GRAF1 forms a complex with MICAL-L1 and EHD1 to cooperate in tubular recycling endosome vesiculation. Front. Cell Dev. Biol. 2, 22 (2014).

98 Sharma, M., Giridharan, S. S. P., Rahajeng, J., Naslavsky, N. & Caplan, S. MICAL-L1 links EHD1 to tubular recycling endosomes and regulates receptor recycling. Mol. Biol. Cell 20, 5181–5194 (2009).

99 Braun, A. et al. EHD proteins associate with syndapin I and II and such interactions play a crucial role in endosomal recycling. Mol. Biol. Cell 16, 3642–3658 (2005).

100 Giridharan, S. S. P., Cai, B., Vitale, N., Naslavsky, N. & Caplan, S. Cooperation of MICAL-L1, syndapin2, and phosphatidic acid in tubular recycling endosome biogenesis. Mol. Biol. Cell 24, 1776–1790 (2013).

101 Xie, S., Farmer, T., Naslavsky, N. & Caplan, S. MICAL-L1 coordinates ciliogenesis by recruiting EHD1 to the primary cilium. J. Cell Sci. 132, jcs233973 (2019).

102 Saha, I., Insinna, C. & Westlake, C. J. Rab11-Rab8 cascade dynamics in primary cilia and membrane tubules. Cell Rep. 43, 114955 (2024).

103 Asante, D. et al. A role for the Golgi matrix protein giantin in ciliogenesis through control of the localization of dynein-2. J. Cell Sci. 126, 5189–5197 (2013).

104 Bergen, D. J. M., Stevenson, N. L., Skinner, R. E. H., Stephens, D. J. & Hammond, C. L. The Golgi matrix protein giantin is required for normal cilia function in zebrafish. Biol. Open 6, 1180–1189 (2017).

105 Follit, J. A. et al. The golgin GMAP210/TRIP11 anchors IFT20 to the Golgi complex. PLoS Genet. 4, e1000315 (2008).

106 Follit, J. A., Tuft, R. A., Fogarty, K. E. & Pazour, G. J. The intraflagellar transport protein IFT20 is associated with the Golgi complex and is required for cilia assembly. Mol. Biol. Cell 17, 3781–3792 (2006).

107 Cockett, M. I., Ochalski, R., Benwell, K., Franco, R. & Wardwell-Swanson, J. Simultaneous expression of multi-subunit proteins in mammalian cells using a convenient set of mammalian cell expression vectors. Biotechniques 23, 402–407 (1997).

108 Rueden, C. T. et al. ImageJ2: ImageJ for the next generation of scienific image data. BMC Bioinformatics 18, 529 (2017).

109 Schindelin, J. et al. Fiji - on Open Source platform for biological image analysis. Nat. Methods 9, 676–682 (2012).

110 Mastronarde, D. N. & Held, S. R. Automated tilt series alignment and tomographic reconstruction in IMOD. J. Struct. Biol. 197, 102–113 (2017).

111 de Chaumont, F. et al. Icy: An open bioimage informatics platform for extended reproducible research. Nat. Methods 9, 690–696 (2012).

