## Supplemental figures for "GRAF1-dependent endocytotic processes and the Golgi apparatus contribute to novel intermediate stages of early ciliogenesis"

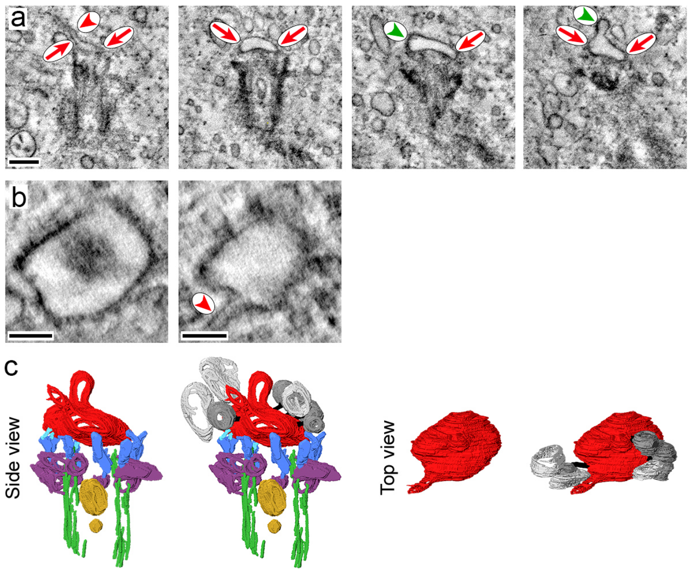


**Suppl. Figure 1. Clathrin-coated buds and incoming vesicles at nascent primary cilia.** (a) Tomogram of a RPE1 cell stably producing centrin1-EGFP and mCherry-Rab8a in which Rab8a accumulated in a punctate pattern at the centrosome after 3 hours of serum starvation. The slices lie ~55 nm apart, red arrows point to the cap-shaped membrane of the nascent cilium, the red arrow head to a tubular extension. Clathrin-coated buds are indicated by green arrow heads. (b) Cross sections of the proximal (left) and distal (right) part of the ciliary membrane structure shown in a. The red arrow head marks a tubular extension. (c) Segmented nascent cilium without (left) and with (right) incoming vesicles. The dark grey vesicular structures are tethered to the nascent cilium, the light grey vesicular structures directly touch it. Microtubules, green; subdistal appendages, purple; distal appendages, dark blue; tethers between distal appendage and nascent ciliary membrane, light blue; nascent ciliary membrane, red; intracentriolar vesicles, yellow. Bars: 200 nm (a), 100 nm (b).

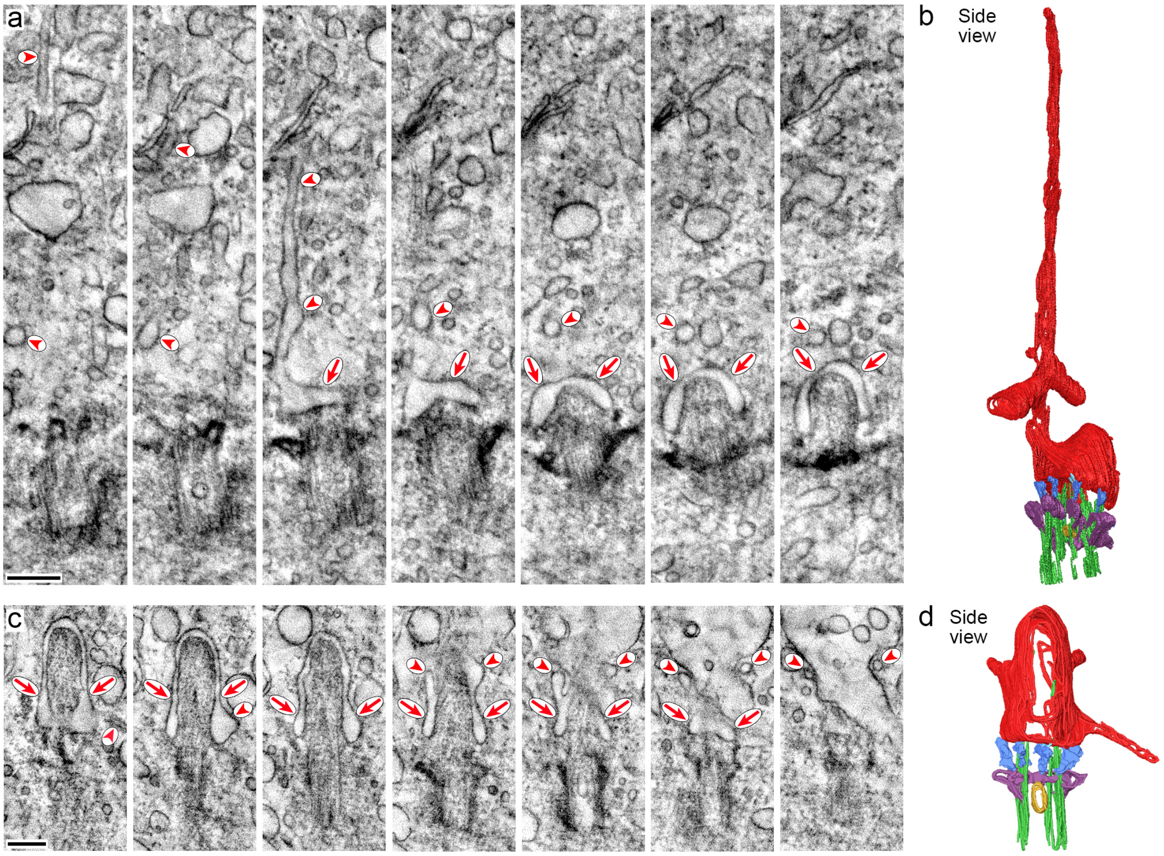


**Suppl. Figure 2. Tubular extensions of nascent primary cilia.** Two tomograms of RPE1 cells stably producing centrin1-EGFP and mCherry-Rab8a in which Rab8a accumulated in a punctate pattern at the centrosome. Cells were serum-starved for 3 hours (a, b) and for 10 hours (b, d), respectively. (a, c) Seven slices lying ~30 nm apart are shown of each tomogram. Red arrows point to the membrane of the nascent cilium, the red arrow heads to tubular extensions. (b, d) The segmented nascent cilia show that extensions of various lengths exist. Microtubules, green; subdistal appendages, purple; distal appendages, dark blue; tethers between distal appendage and nascent ciliary membrane, light blue; nascent ciliary membrane, red; intracentriolar vesicles, yellow. Bars: 200 nm.

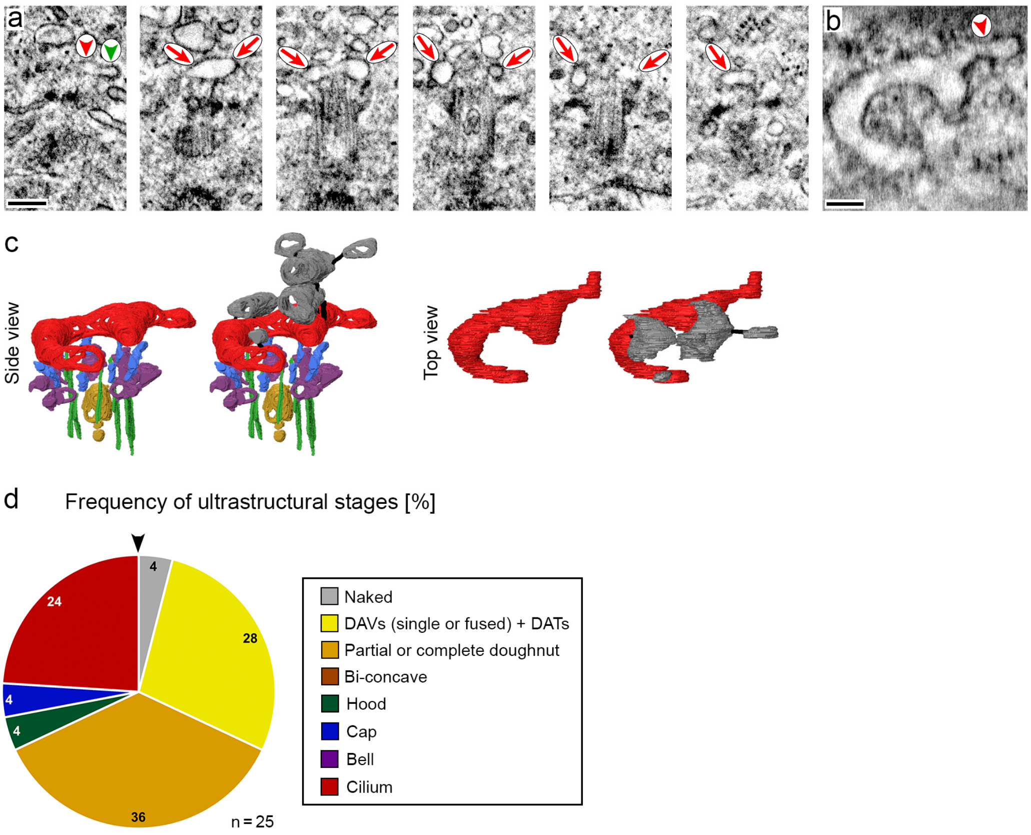


**Suppl. Figure 3. Correlative light and electron microscopy of nascent primary cilia in RPE1 cells producing mCherry.** RPE1 cells stably producing centrin1-EGFP and transiently transfected with an expression plasmid for mCherry were analyzed after serum deprivation for 3.5 and 10 hours. (a-c) Tomogram of a partial doughnut stage in a cell serum starved for 10 hours (tomogram identical to that shown in Fig. 3k, l but rotated around the longitudinal axis). Shown are six slices lying ~55 to 60 nm apart (a), a cross section of the partial doughnut-shaped membrane structure (b) and the segmented nascent cilium (c) without (left) and with (right) incoming vesicles. Red arrows in the tomograms point to the partial doughnut-shaped membrane structure, the red arrow head to a tubular extension and the green arrow head to a clathrin-coated bud. The vesicles in dark grey are attached to the partial doughnut-shaped membrane structure and to each other by tethers (shown in black). (d) Pie chart depicting the frequency (in %) of ciliogenesis stages in cells producing mCherry. Early to late stages are portrayed clockwise starting at the arrow head. *n* indicates the number of tomograms.

Microtubules, green; subdistal appendages, purple; distal appendages, dark blue; nascent ciliary membrane, red; intracentriolar vesicles, yellow. Bars: 200 nm (a), 100 nm (b).

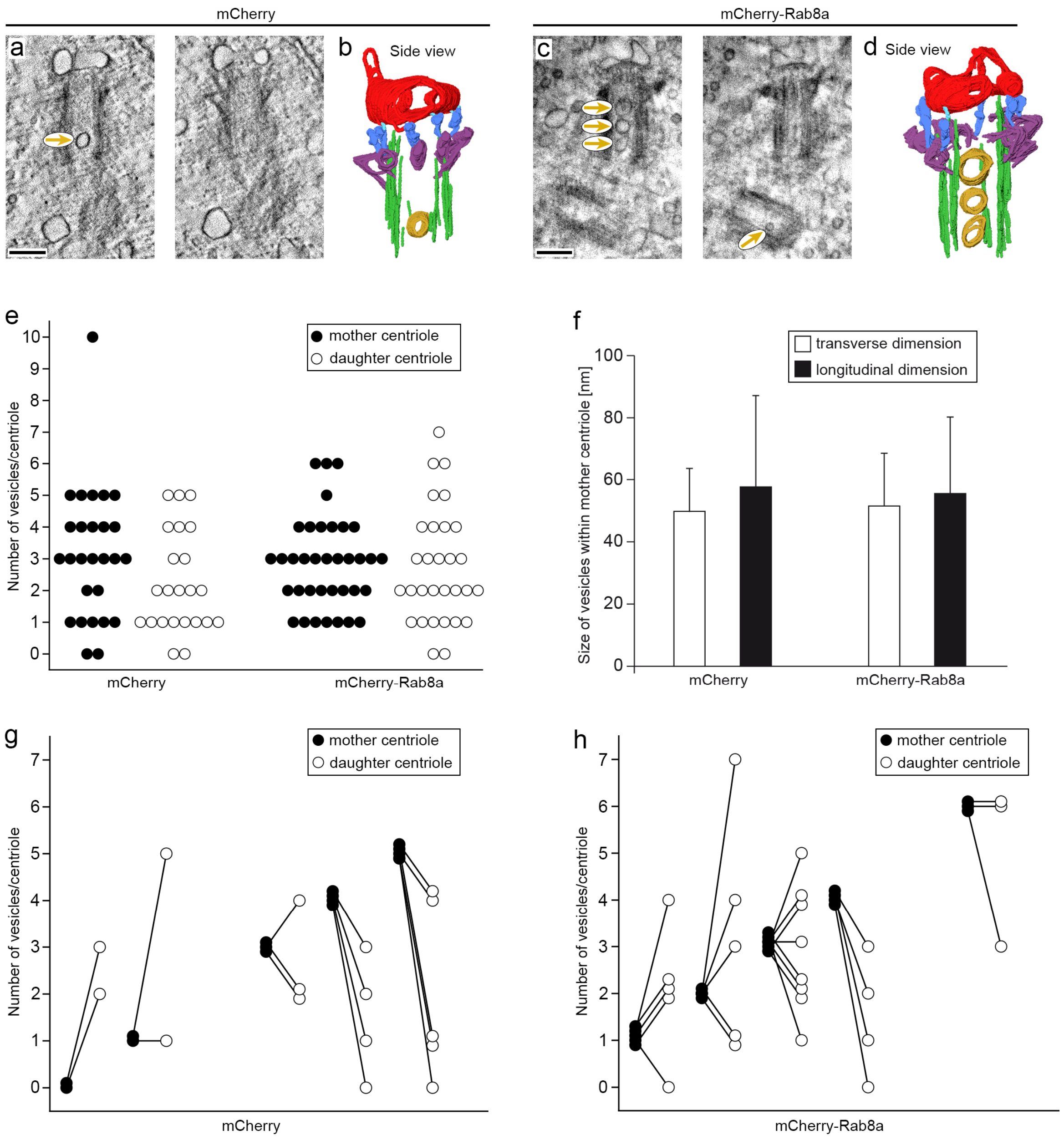


**Suppl. Figure 4. Characterization of intracentriolar vesicles.** (a-d) The centrosomal regions of RPE1 cells producing centrin1-EGFP and either mCherry (a, b) or mCherry-Rab8a (c, d) were subjected to STEM tomography. Cells were serum starved for 10 hours. Two planes in each case are shown which lie ~27 nm (a) and ~40 nm (c) apart. Yellow arrows point to intracentriolar vesicles in mother and daughter centrioles. The segmented nascent cilia can be seen in b and d. (e) Vesicle numbers in mother and daughter centrioles of cells producing mCherry or a mCherry-Rab8a fusion protein. Each circle represents a centriole. (f) Dimension of vesicles inside the mother centrioles in cells producing mCherry or a mCherry-Rab8a fusion protein. The maximum diameter of each vesicle was determined both in the proximal-distal axis (longitudinal dimension) and perpendicular to it (transverse dimension). Shown are the mean + standard deviation. *n* = 85 vesicles (mCherry) and 108 vesicles (mCherry-Rab8a). (g, h) Corresponding number of vesicles between mother and daughter centrioles in cells producing mCherry or a mCherry-Rab8a fusion protein. Microtubules, green; subdistal appendages, purple; distal appendages, dark blue; tethers between distal appendage and nascent ciliary membrane, light blue; nascent ciliary membrane, red; intracentriolar vesicles, yellow. Bars: 200 nm.

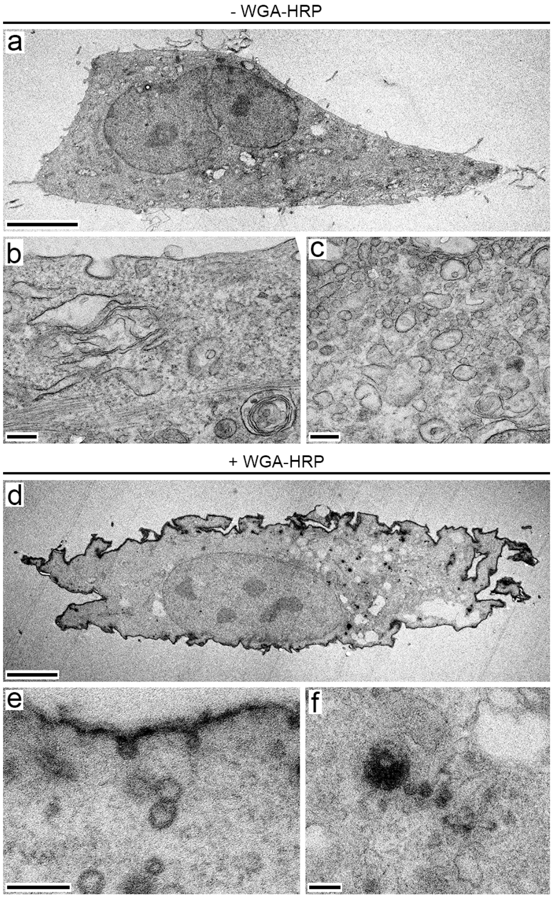


**Suppl. Figure 5. WGA-HRP labeling.** RPE1 cells were incubated in the absence (a-c) or presence (d-f) of WGA-HRP. The overviews (a, d) and higher magnifications of plasma membrane regions (b, e) and of the perinuclear cytoplasm (c, f) demonstrates the specific deposition of electron dense material at the plasma membrane and at intracellular vesicles only when WGA-HRP was present. Bars: 5 µm (a, d), 200 nm (b, c, e, f).
